# Glucose-6-phosphatase is required for organelle reorganization, energy metabolism and motility of *Drosophila* sperm

**DOI:** 10.1101/2024.12.02.626170

**Authors:** Sheida Hedjazi, Tetsuya Miyamoto, Ji-Eun Ahn, Raquel M. Sitcheran, Hubert Amrein

## Abstract

Glucose-6-Phosphatase (G6Pase), a key enzyme in gluconeogenesis and glycogenolysis in the mammalian liver and kidney, converts glucose-6-phosphate to glucose for maintaining systemic blood glucose homeostasis during nutrient deprivation. However, its function has remained elusive in insects, which have no need for G6Pase in sugar homeostasis since they convert glucose-6-phosphate to trehalose, their main circulating sugar, via trehalose phosphate synthase (TPS1). In this study we identify an unexpected and essential requirement for G6Pase in *Drosophila* male fertility, specifically to produce motile sperm. In *G6P* mutant males, spermatogenesis and spermiogenesis appear to proceed normally, leading to the production and transfer of mature sperm. However, once inside the female reproductive tract, *G6P* mutant sperm exhibit severely reduced tail beat frequency and only rarely enter an egg. Moreover, when compared to wild type sperm, *G6P*-deficient sperm are depleted more rapidly from the spermatheca and seminal receptacle, the female sperm storage organs. Immunohistochemical analyses show that *G6P* mutant spermatocytes present with an enlarged and stressed endoplasmic reticulum (ER) and a diminished Golgi apparatus. Additionally, the acrosome, a Golgi derived organelle that is critical for sperm capacitation, exhibits diminished expression of the transmembrane protein SNEAKY, which is essential to breakdown the sperm plasma membrane after fertilization. Metabolic analyses show impairment of both basal and compensatory glycolysis, as well as ATP production, in testes of *G6P* mutant males. Taken together, our investigations unveil a novel and crucial function for G6Pase in male fertility, highlighting its importance in regulating energy homeostasis in reproductive tissues.

## INTRODUCTION

Meiosis is a fundamental process in virtually all metazoans, playing a critical role in the survival of sexually reproducing species, providing a means to increase genetic diversity, and facilitating adaptation to environmental changes over time through natural selection. Genetic screens in *Drosophila melanogaster* have identified hundreds of genes that when mutated cause male sterility. Based on saturation screens, it is estimated that the second chromosome covering roughly 40% of the genome harbors more than 250 different genes essential for male fertility (Wakimoto et al., 2004). Many more genes are likely to be necessary for male fertility, but their function cannot easily be assessed due to their essential roles during development.

*Drosophila* has served as the main invertebrate model system for identifying genetic components essential for male gametogenesis (spermatogenesis), a complex process during which germ cells undergo extensive morphological changes, while also transitioning from diploid germline stem cells (GSCs) to haploid sperm (Figure 1A). In the male fruit fly, approximately 60 to 100 GSCs reside in the apical tip of each testis. Each GSC divides asymmetrically, producing a daughter stem cell and a gonioblast (Dansereau and Lasko, 2008; Fabian and Brill, 2012; Fuller, 1998). The gonioblast, surrounded by two cyst cells that provide support throughout meiosis, first undergoes four mitoses, producing 16 interconnected spermatogonia cells. These 16 cells grow extensively to become primary spermatocytes, before they synchronously enter meiosis to first form 32 secondary spermatocytes and eventually 64 spermatids, which differentiate into mature sperm (Figure 1A). During mating, sperm are transferred to the female reproductive organs, where they are stored in the spermatheca and seminal receptacle until fertilization. Oocytes are fertilized as they pass from the oviduct to the bursa, and the eggs are then laid by the female on suitable substrates that provide nutrients to the larvae that hatch after 24 hours of embryonic development. Sperm stored in the female reproductive organs are viable and capable of fertilization for up to 20 days after mating, and their long-term viability is thought to dependent on nutritional sources provided by the female reproductive organs and the male seminal fluid that is transferred during the mating (Loppin et al., 2015).

**Figure 1:**
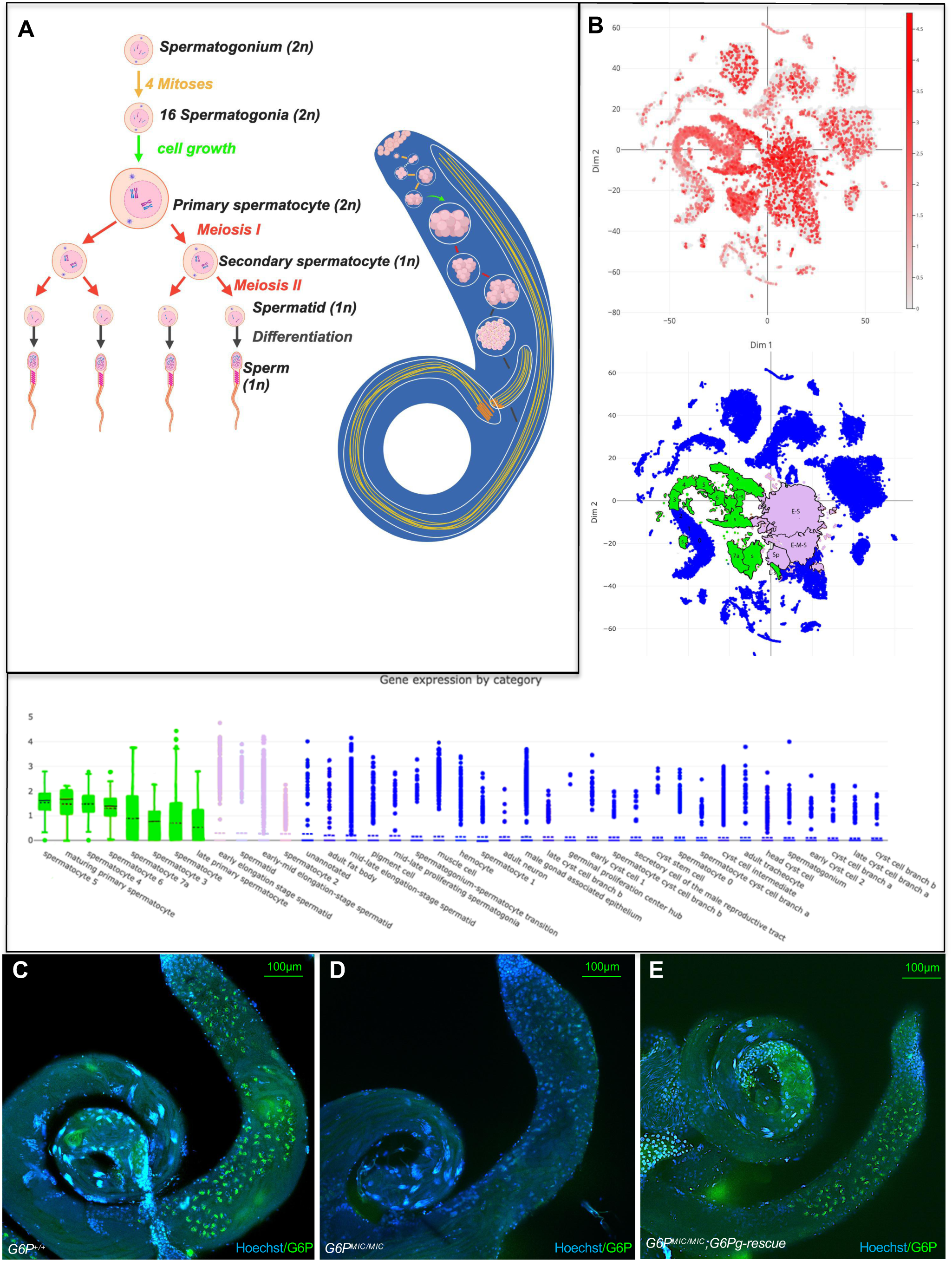
Expression of G6Pase in spermatogenesis. A) Diagram of Spermatogenesis and Spermiogenesis: in Males have approximately 60 to 100 undifferentiated germline stem cells (GSCs) located at the apex of each testis. GSCs divide and produce a primary spermatogonial cell (spermatognium) and another GSC. Spermatogonia undergo four synchronized mitoses, each producing 16 secondary spermatogonia, which grow into larger primary spermatocytes. These spermatocytes undergo synchronized meiosis, generating 64 spermatids, which enter spermiogenesis and transform into mature sperm. Late spermatocytes and spermatids undergo extensive organelle reorganization that is necessary to differentiate into highly specialized sperm. B) Single cell *G6P* RNA expression analysis derived from the “Fly Cell Atlas” (Li et al., 2022). *G6P* mRNA is abundantly present in spermatocytes, but absent in Late elongated spermatids, mature sperm or somatic testis tissue. Tissues/cells are positioned from high (left most) to low (right most) in the graph. Spermatocyte and spermatid cell groups with highest level of expression are shown in green and pink, respectively, while all other cell/tissue types are shown in blue. C-E) G6Pase protein expression in wild type males (C), *G6P* mutant males (D) and *G6P* mutant males containing a genomic rescue construct (E) is visualized using immunostaining with a polyclonal antibody against G6Pase. Immunoreactivity (green) is observed in spermatocytes of wild type and rescue males, but not in *G6P* mutant males. Note that G6Pase is not detected in germ stem cells and spermatids. Hoechst staining was used to visualize DNA/nuclei.

Glucose-6-Phosphatase (G6Pase) is best known for its central role in hepatic gluconeogenesis and glycogenolysis. During times of nutrient deprivation, these biochemical processes can be upregulated, in which G6Pase is the terminal enzyme that hydrolyzes glucose-6-phospate (g-6-p) to glucose, which is then secreted and allows mammals to maintain blood glucose homeostasis. However, the function of G6Pase has remained enigmatic in insects, including *Drosophila*, which do not need G6Pase to generate glucose since they utilize trehalose as their primary circulating sugar. Specifically, when insects break down glycogen (glycogenolysis) or generate trehalose from amino acids (trehaloneogenesis, as opposed to gluconeogenesis), they convert g-6-p to trehalose using trehalose phosphate synthase (TPS) (Matsuda et al., 2015). Surprisingly however, many dipteran insects have maintained a *G6P* gene in their genome (Miyamoto and Amrein, 2019, 2017), and we recently reported that *G6P* has a novel, non-canonical function to facilitate neuropeptide release from discrete populations of *Drosophila* peptidergic neurons the CNS (Miyamoto et al., 2024). During these investigations, we discovered that *G6P* homozygous mutant males are sterile, despite exhibiting normal courtship behavior and mating patterns, suggesting yet another non-canonical function for G6Pase in *Drosophila* spermatogenesis.

In this paper, we show that G6Pase is a metabolic regulator during spermatogenesis. We find that sterile *G6P* mutant males exhibit no deficit in courtship and mate with the same vigor as wild type control males, implying that peptidergic neurons expressing G6Pase are not the cause of male sterility due to aberrant male mating behavior. Instead, we find that the G6Pase enzyme is expressed in spermatocytes during spermatogenesis, consistent with single cell RNAseq analysis (Li et al., 2022). While sperm differentiation appears to proceed normally and *G6P* mutant males transfer sperm to females during mating just like wild type males, tail beat frequency of *G6P* mutant sperm stored in females is severely diminished, and such sperm is depleted sooner from the female reproductive tract than wild type sperm. Cell biological investigations reveal that the endoplasmic reticulum (ER) of *G6P* mutant spermatocytes is more expansive and exhibits increased ER stress, while the Golgi is diminished compared to wild type spermatocytes. Moreover, we find that the acrosome of *G6P* mutant sperm presents with reduced amounts of the cell membrane protein Sneaky, which is necessary for sperm capacitation and essential for the breakdown of the sperm plasma membrane after fertilization. Lastly, we show that testes of *G6P* mutant males have severely impaired glycolytic metabolism and ATP production when compared to wild type testes. Based on these observations and the phenotypic similarities to *G6PC3* mutations in neutrophils of mammals, we propose a model in which *Drosophila* G6Pase functions a metabolite repair enzyme necessary to remove a non-canonical metabolite, and that its accumulation in *G6P* mutant spermatocytes interferes with the cellular transformation of spermatocytes, ultimately leading to deficits in glycolysis and ATP production critical for the high energy demands of sperm.

## RESULTS

### G6Pase is expressed in male germ cells and required for male fertility

While constructing strains to characterize *G6P*’s function in the CNS, we noticed that wild type females failed to produce progeny when mated to homozygous *G6P* mutant males, while homozygous *G6P* mutant females were fully fertile when mated to wild type males. Because *G6P-GAL4* is expressed in diverse subsets of neurosecretory cells in the brain and thoracic ganglion, including cells producing NPF, PDF, Orcokinin, FMRFamide and Nplp 1, and having established that G6Pase facilitates NP release from at least some of these neurons (Miyamoto et al., 2024), we wondered whether impaired male sterility of *G6P* mutants was associated with impaired male courtship and mating behaviors that are under control of some of these neurons. Thus, we mated single *Y/yw; G6P^MIC^/G6P^MIC^* males, *Y/yw* control males and *G6P^MIC^/G6P^MIC^* males carrying a genomic rescue construct (*Y/yw; G6P^MIC^/G6P^MIC^; G6P_rescue/+*) with *G6P* wild type females and measured mating latency and mating time (Figure 2A). By these criteria, *G6P* mutant males and wild type control males were equally efficient in courtship and mating. We next quantified male fertility by measuring oviposition rate and hatching rate of eggs laid by females mated to the three types of males (Figure 2B, Supplementary Table 1). Regardless of the male genotype, females laid about the same number of eggs, suggesting that the seminal fluid, which contains peptides that are essential for increasing production and deposition of eggs after mating (Wigby et al., 2020), is not affected by the loss of *G6P.* However, less than 2% of eggs laid by females mated to *G6P* mutant males developed, compared to more than 80% when mated to control males, while introducing a genomic *G6P* transgene into *G6P* mutants fully rescued fertility (Figure 2B and Supplementary Table 1). Together, these findings show that *G6P* is required for male fertility that is independent of mating behavior and seminal fluid peptide secretion that stimulates egg production.

**Figure 2:**
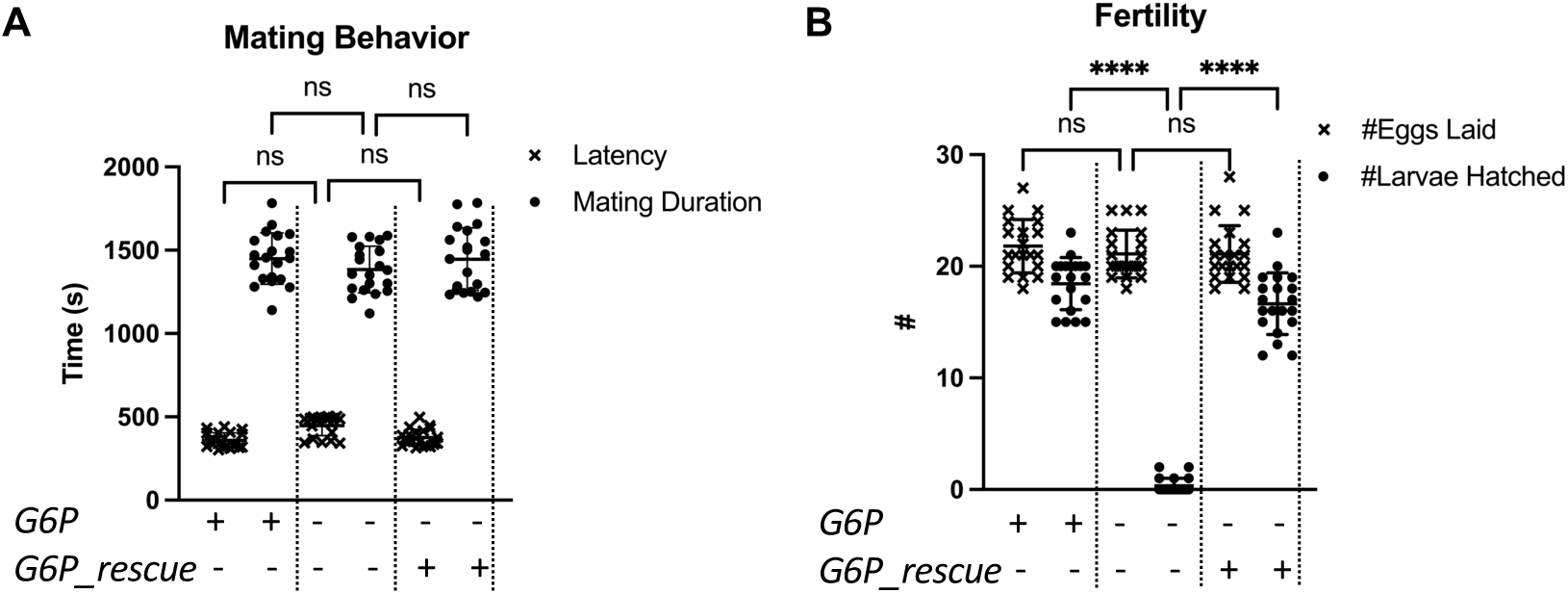
Sterile *G6P* mutant males exhibit normal mating behavior. A) Male courtship was quantified by assessing latency time (x; time from introducing the male and female into the mating arena to successful mounting) and duration of mating (filled circles). No difference was observed between *G6P* mutant and wild type control males, or mutant males carrying a genomic transgene (*G6P_rescue*), mated to mated to *yw* virgin females. n=20. B) Females mated to *G6P* mutant and wild type control males lay the same number of eggs/female per day. Less than 2% of larvae hatched from eggs sired by *G6P* mutant males, whereas approximately 85 % of larvae hatched from eggs sired by control males. Fertility is rescued fully in *G6P* mutant males carrying a genomic *G6P* rescue construct. n=20. Asterisks indicate **** p<0.0001, using two-way ANOVA with the Tukey post hoc test. Error bars represent SE. See also Supplementary table 1.

Single cell transcriptome analysis (Li et al., 2022) indicates that *G6P* is expressed in mid to late spermatocytes and early spermatids (Figure 1B). To confirm that G6Pase is expressed in the male germline, we generated an antibody against a G6Pase peptide (see Methods) and carried out immunostaining experiments of testes from wild type, *G6P* mutant and *G6P* rescue males (Figure 1C-1E). These experiments demonstrate that G6Pase is present in wild type spermatocytes. However, no *G6Pase* immunoreactivity is observed in later stages of spermatogenesis or in somatic cells of testes (Figure 1C). Importantly, no staining is observed in *G6P* mutant males, while staining was restored in spermatogonia and spermatocytes of fertile *G6P* mutant males carrying a genomic *G6P* transgene (Figure 1D and 1E). Of note, inspection of testes from *G6P* mutant males revealed no obvious defect in spermiogenesis, with accumulation of spermatid and mature sperm similar to what was observed in wild type males (Figure 1C-1E and Supplementary Figure 1). Together, these data indicate that *G6P* is expressed in spermatocytes and has an essential role druing spermatogenesis.

### *G6P* mutant sperm are transferred efficiently during mating, but exhibit impaired movement and survival in the female reproductive tract

We next examined whether *G6P* mutant males exhibit phenotypes at later stages in spermatogenesis, including sperm production, sperm transfer and motility, and survival in the female reproductive organs. We introduced *don juan-GFP* (*dj-GFP*), a flagellar, green fluorescent protein-tagged marker that localizes to the sperm tail (Santel et al., 1997), into the *G6P* mutant strain, which permits quantitative visualization of mature sperm in males and females after mating (Figure 3). *dj-GFP* labelled sperm is abundant in seminal vesicle of wild type and *G6P* mutant males (Figure 3B and 3C), with the latter showing a very slight reduction in sperm counts compared to wild type (Figure 3D-F). However, both wild type and *G6P* mutant males transfer *dj-GFP* sperm efficiently during mating (Figure 3H-3I). When we examined the eggs for the presence of *dj-GFP* sperm, we found that 85% of eggs laid by females mated to wild type males were GFP-positive, while only 8 % of eggs laid by females mated to *G6P* mutant males showed GFP signal (Figure 3J). However, in contrast to eggs fertilized by wild type sperm, the eggs fertilized by *G6P* mutant sperm fail to develop, and DAPI staining identifies an elongated nucleus typical of that of sperm cells, suggesting a failure in the breakdown of the sperm cell membrane (Supplementary Figure 2). Together, these observations indicate that mating and sperm transfer occurs normally in *G6P* mutant males, but most of their sperm is unable to enter an egg.

**Figure 3:**
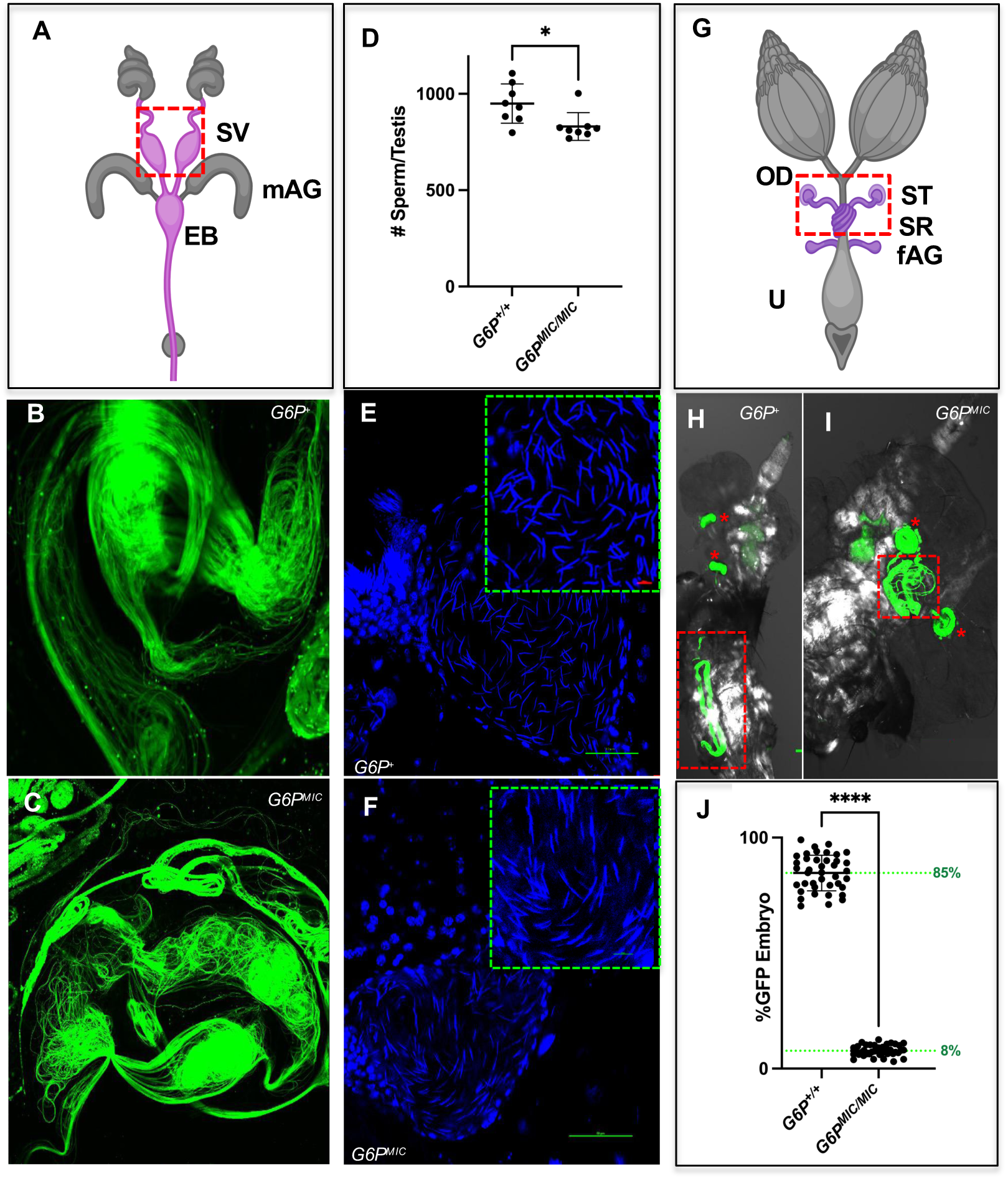
*G6P* mutant males produces and transfer sperm to females during mating. A-C) Sperm storage in males: (A) shows a diagram of the male reproductive system: T=testis, SV=seminal vesicle, mAG: male accessory gland, EB: ejaculatory bulb. Visualization of sperm carrying a dj-GFP transgene in wild type (B) and *G6P* mutant males (C) shows stored sperm in the seminal vesicle. D-F) Sperm present in testis of wild type (E) and *G6P* mutant (F) males. Mature sperm in the seminal vesicle were stained with Hoechst, which stains the elongated sperm nucleus. Sperm count (D) shows a slight reduction of in *G6P* mutants. n = 8 seminal vesicles, asterisks indicate * p<0.05, using unpaired Mann-Whitney U test, error bars represent SE. G-I) Sperm transfer to females: (G) shows organization of the female reproductive system: OD = oviduct, U = uterus, ST = spermathecae, SR = seminal receptacle, fAG = female accessory gland); Wild type female reproductive organs (H) inseminated by sperm from control (*Y/yw; G6P^+^/G6P^+^; dj-GFP/dj-GFP*; left) or (I) *G6P* mutant males (*Y/yw; G6P^MIC^/G6P^MIC^; dj-GFP*/*dj-GFP*; right) is efficiently transferred to females and stored in spermatheca (*) and the seminal receptacle (red squares). J) Sperm from homozygous *G6P* mutant males (*Y/yw; G6P^MIC^/G6P^MIC^; dj-GFP/ dj-GFP*) rarely enter the egg laid by mated females after successful mating, while sperm from control males (*Y/yw; G6P^+^/G6P^+^; dj-GFP/dj-GFP*) do so very efficiently. n = 40 eggs, asterisks indicates **** p<0.0001, using unpaired Mann-Whitney U-test, error bars represent SE.

### *G6P* mutant sperm have severely reduced tailbeat frequency

Key factors for successful egg penetration and fertilization are sperm motility and long-term survival after transfer to the female reproductive tract. Fertilization requires the sperm to sense and move towards the micropyle located between the dorsolateral appendages at the anterior tip of the egg. Given that sperm transfer is not impeded during mating to *G6P* mutant males, but *G6P* mutant sperm are severely compromised in entering the egg (Figure 3D and 3J, respectively), we tested whether sperm motility and survival were affected. We first measured tail beat frequency of wild type and *G6P* mutant sperm in the male (seminal vesicle), which revealed a vigorous tail beat activity for both types of sperm, albeit *G6P* mutant sperm had a small but significant reduction (Figure 4A). Tail beat frequency in the spermathecae of females measured immediately after mating remained similar for wild type sperm, but was dramatically reduced in *G6P* mutant sperm, from 20 Hz to about 3 Hz (Figure 4B). Both phenotypes were rescued by the introduction of a *G6P* genomic transgene into *G6P* mutant males. Moreover, long term storage and/or survival of *G6P* mutant sperm in spermathecae and seminal receptacles was significantly reduced. Specifically, abundance of *G6P* mutant sperm was already reduced after 5 days and almost completely depleted after 10 days in spermathecae, and 15 days after fertilization, neither spermathecae nor seminal receptacle contained any sperm (Figure 3C and 3D). In contrast, wild type sperm was still abundantly present in the seminal receptacle beyond 15 days after mating and remained visible for at least 10 days in spermathecae. Taken together, these observations indicate that *G6P* mutant males produce and transfer sperm in abundance during mating. However, once in the female reproductive tract, *G6P* mutant sperm rapidly lose motility, fail to enter the egg and are more rapidly depleted from the female storage organs when compared to wild type sperm.

**Figure 4:**
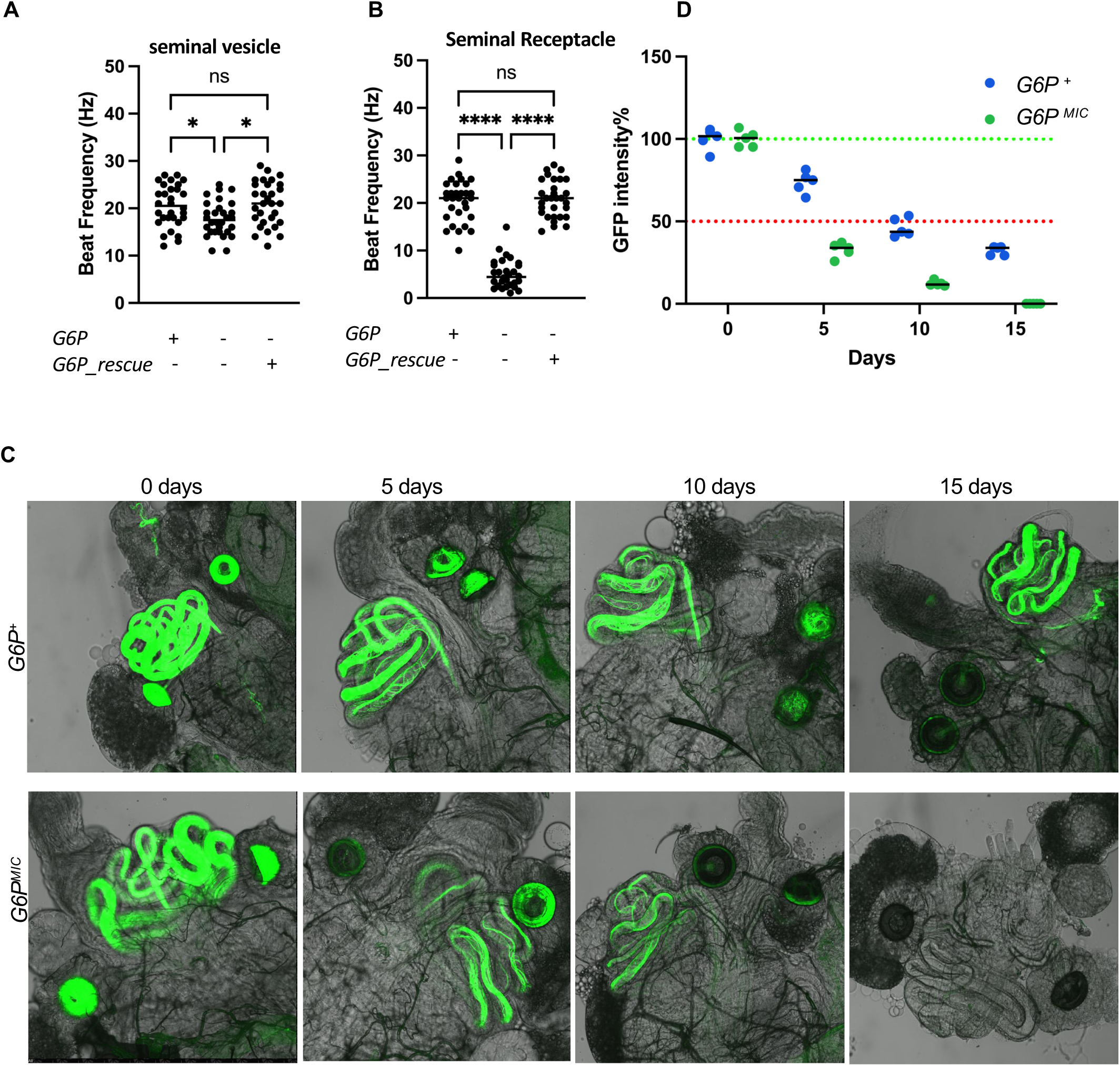
*G6P* sperm exhibit severely reduced sperm tail beat and survival. A-B) Tail beat frequency of wild type and *G6P* mutant sperm dissected from male seminal vesicle before mating (A) or dissected from seminal receptacle (B) n =30, asterisks indicate* p<0.05, and **** p<0.0001, using one-way ANOVA with the Tukey post hoc test. error bars represent SE. C) Survival of control sperm (top line, *Y or y w; G6P^+^; dj-GFP*) and *G6P* mutant sperm (bottom line, *Y or y w; G6P^MIC^; dj-GFP*) in the female reproductive organs after mating (0 days), or 5, 10 and 15 days after mating. *G6P* mutant sperm is depleted sooner in both sperm storage organs, compared to wild type sperm. D) Total average *GFP* fluorescence derived from sperm of *Y or y w; G6P^+^; dj-GFP* (blue) and *Y or y w; G6P^MIC^; dj-GFP* (green) sperm in female reproductive tract immediately and 5, 10 and 15 days after mating. n=5

### Loss of *G6P* disrupts organelle reorganization during spermatogenesis

*Drosophila* spermatocytes undergo impressive, metabolically demanding growth during meiosis. They increase in size by about 25-fold in the G2 phase of the cell cycle (meiotic prophase), a process that requires substantial metabolic investment resulting in cells that are larger than most somatic cells (Lindsey and Tokuyashu, 1980). Furthermore, extensive, energetically costly reorganization of organelles, such as the mitochondria, endoplasmic reticulum (ER) and Golgi apparatus, is necessary to facilitate the morphological transformation that occurs during spermiogenesis, from large spherical spermatocytes to the elongated soma with and exceedingly long tail (up to 1.8 mm) of mature sperm. During this process, the ER is broken down, mitochondria are relocated to what becomes the mid piece of the sperm where they fuse into the mitochondrial sheet powering the sperm tail during fertilization, and the Golgi compartment constricts and moves to the apical tip to form the acrosome (Fabian and Brill, 2012; Fitch and Wakimoto, 1998; Wilson et al., 2006). Thus, we assessed whether loss of *G6P* had any effect on this cellular reorganization. Consistent with mammalian G6Pases, which reside in the ER (Burchell et al., 1994), we find that in *Drosophila* spermatocytes, G6Pase is perinuclear and co-localizes with the ER resident protein calnexin (Figure 5A). Furthermore, size comparison revealed a significant expansion of the ER in *G6P* mutant spermatocytes compared with wild type (Figure 5B). To assess whether this expansion is associated with ER stress, we examined the effect of *G6P* loss on total calnexin amount. Indeed, total calnexin, which is upregulated when cells are under stress (Coe et al., 2008; Danilczyk et al., 2000; Jackson et al., 1994), was significantly elevated in *G6P* mutant spermatocytes when compared to wild type spermatocytes (Figure 5C). We also quantified splicing of *XBP1* mRNA, an ER stress induced transcriptional regulator which plays a key role in the Unfolded Protein Response (URP) (Cairrão et al., 2022; Uemura et al., 2009). Comparison between mRNA isolated from testes of wild type and *G6P* mutant males revealed that spliced *XBP1* mRNA is significantly increased in *G6P* mutants at the expense of non-spliced *XBP1* RNA (Figure 5D). Additionally, Binding immunoglobulin Protein (BiP), a UPR chaperone (Bertolotti et al., 2000), is also upregulated in *G6P* mutant spermatocytes (Supplementary Figure 3). Together these findings indicate robust induction of an ER stress response in spermatocytes that lack a functional G6Pase.

**Figure 5:**
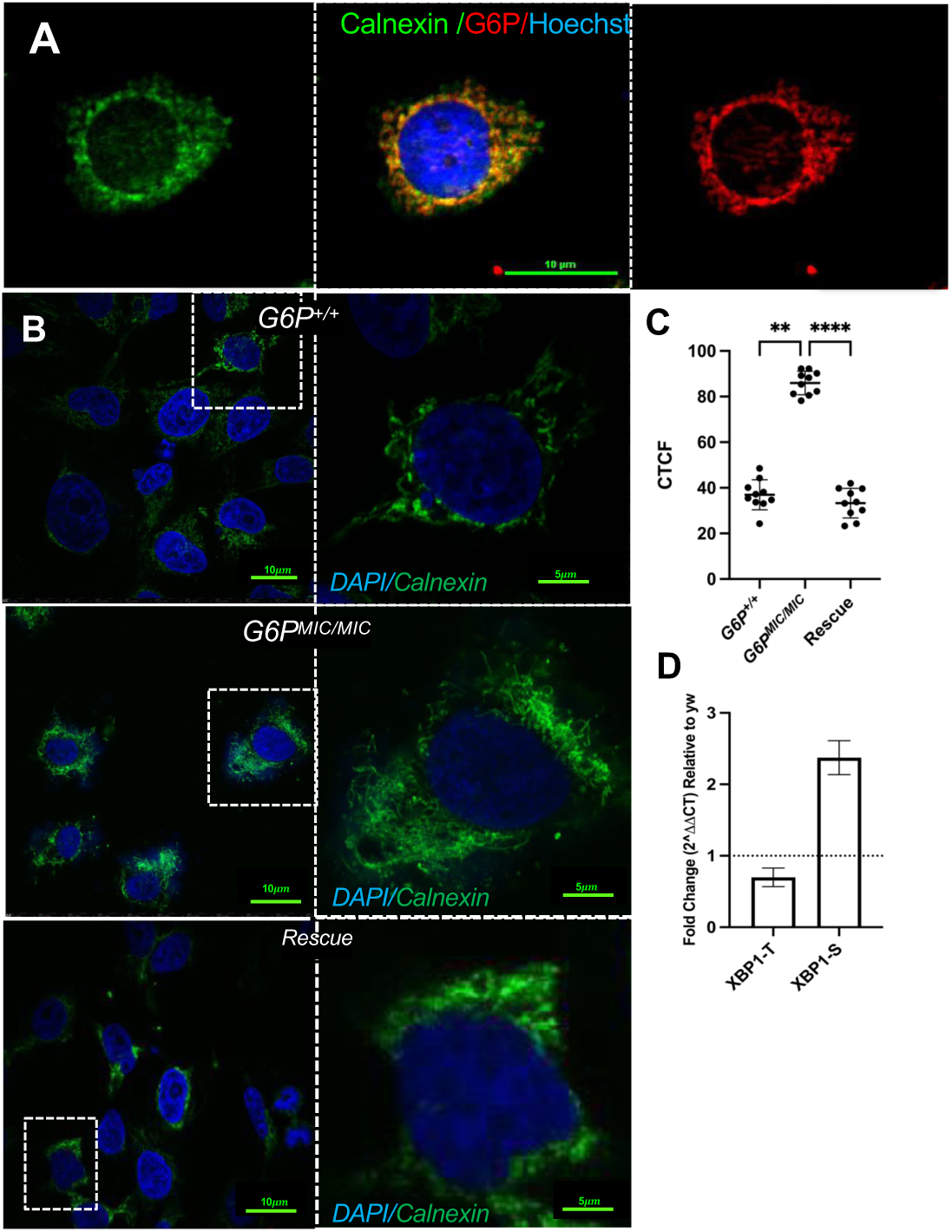
G6Pase protein expression and Endoplasmic Reticulum stress in spermatocytes lacking *G6P*. A) Anti-G6Pase antibody staining reveals perinuclear association with and co-localization with calnexin in the ER. DNA was visualized using Hoechst. B and C) ER is enlarged in *G6P* mutant spermatocytes, with concomitant elevation of calnexin levels. DNA was visualized using Hoechst in B. Quantification of calnexin is shown as Corrected total cell fluorescence (CTCF) using Image J. ** indicates p<0.01, and **** indicates p<0.0001, using one-way ANOVA with the Tukey post hoc test. error bars represent SE. D) *G6P* mutant spermatocytes show ER stress evident from increased splicing of the transcription factor encoding *XBP1* mRNA. Splicing was quantified by RT-PCR using RNA isolated from whole testes of wild type and *G6P* mutant males non-spliced (XBP1-T) and spliced (XBP1-S) was normalized and compared to wild type (dotted line).

We next analyzed the Golgi in wild type and *G6P* mutant spermatocytes using an antibody against the Golgi Matrix protein 130 (GM130) and found *G6P* mutant spermatocytes had a severely reduced number of Golgi compartments (Figure 6A and 6B), similar to peptidergic neurons of *G6P* mutant flies (Miyamoto et al., 2024). The Golgi gives rise to the acrosome, which is thought to be involved in sperm-egg (i.e. micropyle) recognition (Intra et al., 2006; Loppin et al., 2015; Morciano et al., 2021; Perotti and Riva, 1988) and the initiation of development post-fertilization (Fitch and Wakimoto, 1998; Wilson et al., 2006). To assess acrosome integrity, we crossed a *sneaky-GFP* (*snky-GFP*) transgene, which encodes a membrane protein localized to the acrosome (Wilson et al., 2006), into the *G6P* mutant strain. Indeed, the acrosome of *G6P^MIC^; snky-GFP* sperm exhibited less GFP immunofluorescence compared to *G6P^+^; snky-GFP* sperm (Figure 6C and 6D). In summary, *G6P* mutant spermatocytes show an expanded ER exhibiting elevated stress, and a concomitant reduction of the Golgi and partial loss of SNEAKY, an integral transmembrane protein of the acrosome that is essential during and after fertilization (Fitch and Wakimoto, 1998; Wilson et al., 2006).

**Figure 6:**
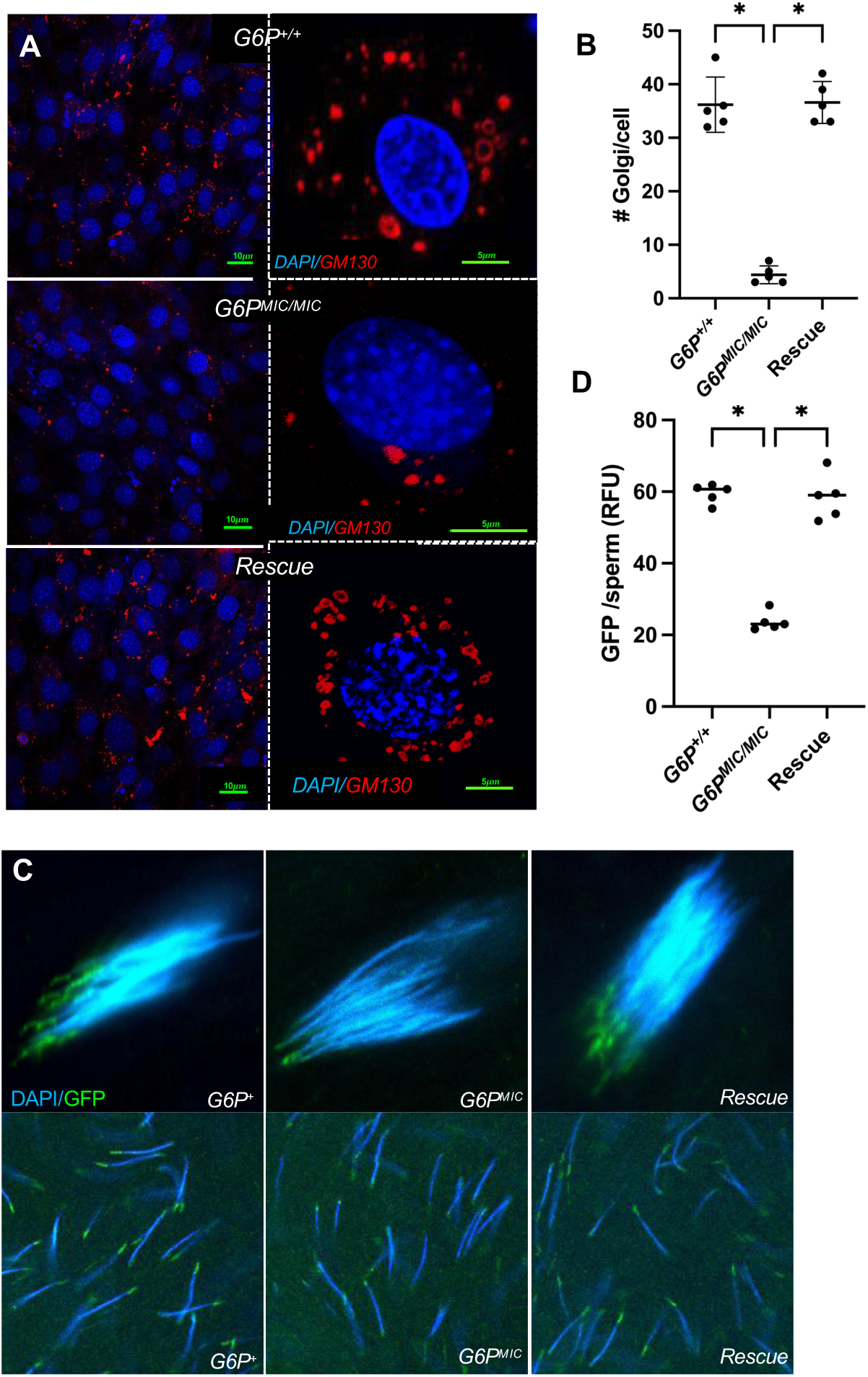
Loss of *G6P* leads to a reduction of Golgi in spermatocytes and the acrosome membrane associated protein SNEAKY. A) *G6P* mutant spermatocytes (top, nuclei stained with Hoechst in blue) show a significant reduction of Golgi compartments, visualized using an antibody against the Golgi marker protein GM130 (red), in spermatocytes compared to wild type spermatocytes (bottom). Images to the right show higher magnification of a single spermatocyte. B) Quantification of the average number of Golgi compartments/spermatocyte, determined by counting the cells and Golgi compartment of representative field image areas (n) as shown in A. n = 5, asterisks indicate * p<0.05, using one-way ANOVA with the Tukey post hoc test, error bars represent SE. C) *G6P* mutant sperm show reduced GFP-tagged SNEAKY protein in the acrosome, compared to wild type sperm. Top image shows sperm bundle in a cyst. Bottom images show free, mature sperm. D) GFP intensity shown as RFU (relative fluorescence units)/sperm measured from representative images of dissected testes (n) using Image J. N = 5, asterisks indicate * p<0.05, using one-way ANOVA with the Tukey post hoc test, error bars represent SE.

### Testes of *G6P* mutant males exhibit reduced energy metabolism

Our studies have shown that *G6P* mutant sperm are incapable of fertilizing eggs due to severely compromised motility and reduced survival in spermathecae and seminal receptacle. The loss of motility suggests that *G6P* mutant sperm have an energy deficit, likely caused by increased ER stress and a diminished Golgi. To investigate potential metabolic dysfunction in *G6P* mutant male testes, we conducted ex vivo energy consumption measurements using a Seahorse XF analyzer (Agilent Technologies). We observed a 60% reduction in both basic and compensatory glycolysis when compared to wild type testes (Figure 7A), demonstrating an impaired ability to upregulate glycolysis when mitochondrial respiration is inhibited. Moreover, G6P mutants exhibit impaired mitochondrial function, resulting in significantly reduced basal respiration, spare respiratory capacity and ATP production (Figure 7B). These observations are consistent with recent *ex vivo* metabolic measurement, which showed that both glycolysis and mitochondrial respiration are major metabolic pathways in *Drosophila* sperm (Wetzker et al., 2024). Reduced ATP levels in testes of *G6P* mutant males were independently confirmed by measuring total ATP levels using the ATPLite system (PerkinElmer), which revealed that *G6P* mutants contained only about 50% of the ATP when compared to testes of wild type or *G6P* mutant males carrying the genomic rescue transgene (Supplementary Figure 4). Collectively, these results imply that sterility of *G6P* mutant males is caused by a cascading series of events, starting with a failure to transform cellular organelles during spermatogenesis and ultimately resulting in multiple metabolic deficiencies that severely impacts sperm motility and survival.

**Figure 7:**
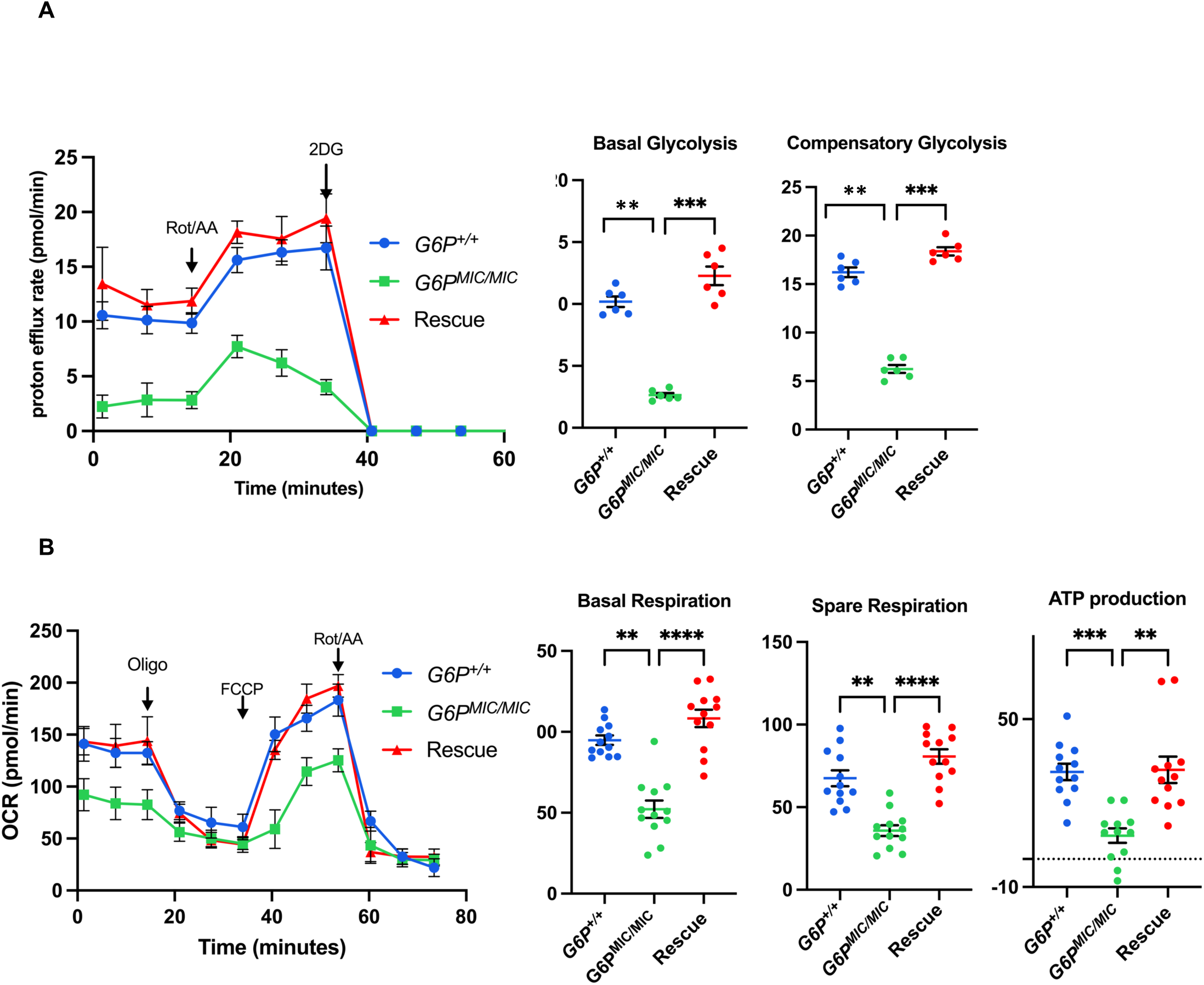
Testes of *G6P* mutant males have severely reduced energy metabolism. Metabolic capacity of testes from wild type (blue) and *G6P^-/-^* (green) males was determined using a Seahorse XF analyzer (Agilent Technologies). *G6P* mutant flies carrying a genomic rescue construct (*G6P^MIC^/G6P^MIC^*; *G6P_rescue*) are shown in red. A) Basal glycolysis is impaired in testes of *G6P* mutant males, and inhibition of the ETC using rotenone/antimycin A shows that compensatory glycolysis is also reduced. B) Overall oxygen consumption is reduced in testes of *G6P* mutant, with significantly diminished basal respiration, spare respiratory capacity and ATP production. Note that *G6P* is not expressed in somatic tissues of testes, suggesting that the germ line probably accounts for the energy deficit. N=5, asterisks indicate** p<0.01, *** p<0.001, and **** p<0.0001, using one-way ANOVA with the Tukey post hoc test. Error bars represent SE.

## DISUCSSION

In this paper, we established a novel function for the metabolic enzyme G6Pase in *Drosophila* male fertility that is distinct from its known role as a regulator of glucose metabolism in mammals. Given our recently reported role for G6pase as a non-canonical facilitator of neuropeptide release in several types of peptidergic neurons in the CNS (Miyamoto et al., 2024), we first considered the possibility that male sterility had a behavioral cause, whereby reduced neuropeptide signaling would interfere with proper male courtship and mating behavior. However, *G6P* mutant males court and mate normally, and they produce and transfer sperm during copulation just like wild type males. Analysis of spermatogenesis established that *G6P* has a cell autonomous function in spermatocytes that is necessary for proper organelle reorganization which is essential to produce capacitated sperm. Specifically, we find that the *G6P* gene is expressed during spermatogenesis and show that G6Pase is present in the ER of spermatocytes and early spermatids. Second, we discovered that *G6P* mutant spermatocytes show ER stress and a compromised Golgi apparatus, and that the acrosome of *G6P* mutant sperm have diminished amounts of the essential acrosome membrane protein SNEAKY. Third, *G6P* mutant sperm are much less motile and vigorous after mating compared to wild type sperm, and their survival in two main female storage organs, the spermathecae and the seminal receptacle, is shortened. And fourth, testis of *G6P* mutant males exhibit a sharply reduced glycolytic and mitochondrial energy metabolism. Overall, these results suggest broader roles for G6Pase as an integrator of metabolic homeostasis in bioenergetically-demanding tissues.

G6Pases have mainly been studied in mammals, particularly in the context of gluconeogenesis and glycogenolysis in the liver and kidney, mediated by *G6PC1*. There, G6Pase converts glucose-6-phosphate to glucose, which is released into the blood stream to ensure systemic glucose homeostasis during times of limited nutrient availability. Patients with mutations in *G6P1C* suffer from glycogen storage disease type I and present with hypoglycemia and hyperlipidemia, and eventually with hepatomegaly due to glycogen accumulation in the liver (Chou et al., 2015). In addition to *G6PC1*, mammals have two additional G6Pase encoding genes. *G6PC2* is mainly expressed in in the pancreas and has been associated with cell-mediated autoimmunity, and patients with a mutation in *G6PC2* and *G6PC2* knock out mice have lower fasting blood glucose levels, due to increased glycolysis in β-cells (Bosma et al., 2020b, 2020a). Lastly, *G6PC3* is broadly expressed in many tissues, but it has mainly been studied in neutrophils, where congenital *G6PC3* deficiency leads to neutropenia due to failure to clear 1,5-anhydroglucitol phosphate (1,5AG6P) from cells (Veiga-da-Cunha et al., 2019). Interestingly, while all three mammalian G6Pase enzymes can hydrolyze g-6-p, their catalytic activities and substrate specificities differ, and consequently, so do their specific roles in metabolism (Veiga-da-Cunha et al., 2019). We note that while *Drosophila* G6Pase can convert amino acids via g-6-p to glucose in peptidergic neurons in the brain (Miyamoto and Amrein, 2019), it remains to be seen whether g-6-p is the main substrate for this enzyme in either peptidergic neurons or spermatocytes.

### Loss of *G6P* in spermatocytes affect pre-and post-fertilization properties of sperm

Regardless of the specific mechanism through which G6Pase functions in spermatocytes and facilitates the reorganization of organelles during spermiogenesis, its loss has severe consequences on sperm, evidenced by the reduced metabolic activity of *G6P* mutant testes and the loss of sperm motility in the female reproductive organs. Of note, while motility of *G6P* mutant sperm is slightly, though significantly, impaired in the seminal vesicle compared to wild type sperm, this deficit is exacerbated when sperm motility was assessed in the seminal receptacle of females (Figures 3A and 3B). Increased loss of motility might reflect continuing exhaustion of available energy as a function of time. Interestingly, Wetzker and colleagues recently showed that there is a pronounced shift in sperm energy production associated with the environment, observing an increase in OXPHOS after sperm transfer to the female reproductive system (Wetzker et al., 2024). It seems likely that external factors in male seminal fluid and excretions from female accessory glands, which mix with sperm during and immediately after copulation, respectively, contribute to this metabolic shift. For example, *G6P* mutant sperm might lack critical membrane components, especially in the acrosome (see below) that are necessary for sperm capacitation, and thus fail to respond to the new external environment encountered in the female reproductive tract, which is known to provide nutrients and signals essential for long-term sperm motility (McCullough et al., 2022; Zelinger et al., 2024). Such defects are quite likely, given that the severely diminished Golgi and deficits of acrosome associated proteins might reflect impaired post translational modification of proteins involved in secretory pathways and glycosylation of sperm proteins.

The acrosome has essential roles during sperm capacitation and fertilization, albeit there are pronounced differences across species. In mammals, the acrosome plays a critical role in the fusion of the sperm and oocyte membranes (Okabe, 2018), while in *Drosophila* and probably other insects, membrane fusion does not occur, as sperm enter an oocyte fully intact through the micropyle. While roles for the *Drosophila* acrosome have been postulated during eggs-sperm interaction (see below), it has been well established that after sperm entry, acrosome components are essential in several processes for successful fertilization and initiation of development. Specifically, in eggs inseminated by sperm lacking the acrosome associated protein SNKY, the male nucleus fails to decondense and form a functional pronucleus, leading to an inability to initiate embryonic development (Fitch and Wakimoto, 1998; Wilson et al., 2006). Interestingly, membrane permeability of *snky* mutant sperm in fertilized eggs is reduced when compared to wild type sperm, implying that the breakdown of the sperm cell membrane requires SNEAKY protein (Fitch and Wakimoto, 1998).

Additional roles for the acrosome during sperm entry are implied by specific glycosylation signature present on the acrosome membrane (Perotti and Pasini, 1995). These signatures appear to be complementary with sugar residues mapped to the micropyle of the oocyte, suggesting possible interactions between the two membranes (Intra et al., 2006; Perotti and Riva, 1988). Evidence that glycosylation is critical for *Drosophila* sperm-egg interaction has been obtained from *Casanova* (*cas*) mutant males, which produce mature motile sperm that are, however, unable to fertilize the egg (Perotti et al., 2001). *Cas* mutant sperm have only half of the β-hexosaminidase activity when compared to wild type sperm, with none of it on the membrane overlaying the acrosome itself. Unfortunately, the *cas* mutant strain has been lost before the gene could be mapped and molecularly characterized. We interrogated single cell expression data in the male reproductive tract for key enzymes of the hexosamine biosynthetic pathway (HBP), which indeed shows that respective genes are expressed in spermatocytes (Supplementary Figure 5), lending further support for a role of membrane glycosylation in sperm capacitation. Thus, it will be interesting to see whether the diminished Golgi of *G6P* mutant spermatocytes affects expression or function of HBP enzymes in a manner similar to neutrophils of humans with mutations in *G6PC3*, which are highly defective in glycosylation (Hayee et al., 2011).

### A non-canonical function for *Drosophila* G6Pase in the male germ line

*Drosophila* G6Pase in spermatogenesis and neurosecretory cells in the CNS is unlikely used for systemic glucose production, as is the case for hepatic G6Pase in mammals. Neurosecretory cells and spermatocyte perform energy consuming tasks – production of large amounts of neuropeptides and neural signaling in the CNS, proliferation, meiosis and cellular and organelle reorganization in the case of spermatogenesis, and hence are not equipped to engage in energy consuming processes of glucose production for the benefit of systemic glucose homeostasis. Moreover, other genes encoding enzymes required for either gluconeogenesis (*e.g.,* 1,6-bisphosphatase and phosphoenolpyruvate carboxykinase) or glycogenesis and glycogenolysis (glycogen synthase, debranching enzyme, glycogen phosphatase) are not expressed in spermatocytes (Supplementary Figure 6)(Li et al., 2022), which further supports a non-canonical function of *Drosophila* G6Pase in the male germ line.

We propose that *G6P* in spermatocytes functions similar to *G6PC3* in neutrophils. Neutrophils lacking *G6PC3* are sensitive to the accumulation of 1,5-Anhdrogucitol (1,5AG), a glucose analog and common monosaccharide present in most foods. 1,5AG enters cells via glucose transporters, where it is phosphorylated by hexokinase to 1,5AG6P. However, 1,5AG6P is non-metabolizable and must be cleared from cells via G6Pase by converting it back to 1,5-Anhdrogucitol (1,5AG), and then exported via glucose transporters and excreted through the kidneys. In neutrophils of patients homozygous for *G6PC3* mutations, 1,5AG6P accumulates and disrupts glycolysis, leading to the well-known cellular phenotypes, including ER stress, reduction of glycolytic metabolites such as lactate, and glycosylation deficits (Hayee et al., 2011; Jun et al., 2011, 2010), ultimately resulting in loss of neutrophils (Veiga-da-Cunha et al., 2019). Of note, *G6PC3* deficiency has also been associated with numerous additional pathologies, including urogenital malformations, heart defects, and learning disabilities (Banka et al., 2011; Boztug and Klein, 2009), suggesting that other tissues and organs are sensitive to the accumulation of 1,5-AG6P, or possibly other non-metabolizable compounds (Dienel, 2020). Thus, G6Pase encoded by *G6PC3* functions as a metabolite repair enzyme, like several other enzymes in the glycolysis pathway (Bommer et al., 2020). Metabolite repair enzymes are necessary to eliminate non-canonical products that occur during glycolysis due to substrate promiscuity inherent to most glycolytic enzymes, including hexokinase (Veiga-da-Cunha et al., 2020). Based on the similarity of the cellular phenotypes between *G6P* mutant spermatocytes and *G6PC3* mutant neutrophils (ER stress, reduced Golgi compartments), and the severe disruption of glycolysis in testes of *G6P* mutant males, we suggest that failure of *G6P* mutant sperm to fertilize eggs is brought about by a non-metabolizable compound that interferes with glycolysis in male germ cells. Of note, the glucose transporter GLUT3 is robustly expressed in spermatocytes and spermatids (Li et al., 2022), and hence, this transporter might participate as a metabolic repair protein, similar to the glucose transporter expressed in human neutrophils (Veiga-da-Cunha et al., 2019). Future studies will be necessary to determine if any and what kind of sugar compounds might be the substrate for *G6P* in *Drosophila* spermatocytes.

Based on the Mammalian Reproductive Genetics Database, testes expression of *G6PC3* in humans and rodents is most abundantly found in spermatocytes and Sertoli cells, which support spermatocytes (https://orit.research.bcm.edu/MRGDv2). Of note, almost half of men carrying *G6PC3* mutations have urogenital defects, with the most prominent being cryptorchidism (failure of testicles to descent during embryonic development)(Yeshayahu et al., 2014). Thus, it will be interesting to uncover the specific metabolic role of *G6PC3* in mammalian testes and germ cells, including those of man, and determine if it plays a role in fertility.

## ACKNOWLEGEMETNS

We thank Drs. Barbara Wakimoto (University of Washington), Mariana Wolfner (Cornell University), Mia Levine (University of Pennsylvania) and Harmit Malik (Fred Hutchinson Cancer Center) for *Drosophila* strains. We thank Drs. Malea Murphy (Integrated Microscopy and Imaging Laboratory) for assistance with fluorescence microscopy and Robbie Moore (Flow Cytometry and Cell Sorting Facility) for help with the Sea horse assay. This work was supported by grants from the National Institute of Health (NIH-1R21NS1118118-01 and NIH-R01 DC018403-01A1) to Hubert Amrein. The authors declare no conflict of interest.

## METHODS

### *Drosophila* Stocks and Husbandry

*Drosophila* stocks were maintained in 12-hour light/dark cycle on standard corn meal food in plastic vials at 25° C. The *y w^1118^* strain was used as wild type control. Fly stocks used were: The *G6P* mutant strain (*G6P^MIC^*) and the *dj-GFP* transgenic strain was generated from the Bloomington stock center (#57904) by back-crossing it *y w^1118^* and removing the *protamineB-eGFP* transgene. The *dj-GFP*/*dj-GFP*; *G6P^MIC^/G6P^MIC^* strain was generated by combing the *G6P^MIC^* allele (chromosome 2) and the *dj-GFP* transgene (chromosome 3), balanced with *GyO* and *TM3 Sb*.

### Molecular Cloning

To generate the genomic *G6P* rescue transgene, a 4,925 base pair long fragment was amplified from genomic DNA including the entire coding sequence with introns, 3,109 base pairs upstream of the ATP start codon and 500 base pairs downstream of the stop codon using a forward primer (5ʹ-GGACTA<u>GCGGCCGC</u>CTACTGCTCCACAAGAAGGAGATGACTT −3ʹ) and a reverse primer (5ʹ-ACCTTA<u>GGTACC</u>TCCCATCATGGCTATCACGCGCTGCGAA −3ʹ). NotI and KpnI (underlined) restriction enzyme sites were incorporated into primers for directional cloning. The PCR product were then subcloned into pCaSpeR4 transformation vector (Addgene). Transgenic fly lines were generated by a standard procedure with P-element mediated transformation of *w^1118^* embryos (Rainbow transgenic Flies Inc., Camarillo, CA).

### Statistical analysis

Statistical significance was assessed using either the Student’s t-test or a two-tailed Mann–Whitney U test for pairwise comparisons. For multiple comparisons, one-way ANOVA was applied with Tukey’s post hoc test or the Kruskal–Wallis test with Dunn’s post hoc test. Error bars represent the standard error of the mean (SEM), unless stated otherwise. Significance levels were denoted as *P < 0.05, **P < 0.01, with NS indicating no significance. All replicates were derived from distinct biological samples (different flies or cells). Image analysis was conducted using the NIS-Elements software suite, and statistical analyses were carried out with Real Statistics Add-Ins in MS Excel and GraphPad PRISM.

### Fertility and mating test

All mating experiments were performed with males 4-7 days of age. Single pair matings were set up using a mating wheel. Naive males (collected and isolated within 24 hours after hatching) were kept in isolation for at least 3 days on standard food before used in mating assays. A single male was paired with a single virgin female (2 to 5 days old) in a mating chamber of the mating wheel. The latency time is defined as the time from the moment the male and female were introduced into the mating chamber until the male mounted the female. Mating or copulation time is defined as the time interval from mounting to dismounting the female. To test for fertility, three females were crossed with three males for 48 h on a grape juice plate with yeast paste. After day 2, the males were removed from the chamber, and each female was transferred to a fresh grape juice plate with yeast paste. The number of eggs was counted after 24 h. The plates were then placed in a humidity chamber for the next 72 h, and the number of hatched larvae was recorded.

### Immunostaining

Fly testes dissection, fixation, and staining were performed as described in Sitaram et.al. (Cytological analysis of spermatogenesis: live and fixed preparations of Drosophila testes., 2014) with minor modifications. For the intact testes, a hydrophobic barrier pen (Super PAP Pen,71310, EMS) was used to border the testis and protect them from getting squished at the start of the process. 2-3 pairs of testes were dissected in 1x PBS and transferred to the glass microscope slide marked with a PAP pen. A coverslip glass was placed very gently on top of the preparation. For specific stages of cell staining, testes were dissected and transferred to a coverslip containing 20uL of PBS. Testes were cut open to retrieve spermatocytes. A glass slide was gently placed on top of the coverslip, and the excess PBS was removed using kimwipe, with ensuing pressure pushing spermatocytes out. The sample was immediately frozen in liquid nitrogen. Next, the coverslip was removed, and the glass slide was placed in cold 95% ethanol for 10 minutes in −20°C freezer. Samples were transferred to 4% Formaldehyde in PBS with 0.1% triton (PBST 0.1%) for 10 minutes, washed in PBS 3X each time for 5 minutes and then transferred to PBST for 45 minutes to permeabilize the cell membrane. Samples were washed again 3 times for 5 minutes each in PBS and blocked by immersing the slides in PBS with 1% BSA for 45 minutes and then incubated overnight in a 4°C fridge with primary antibody solution diluted in PBS containing 1% BSA. The next day, preparations were washed once with PBST and twice with PBS for 5 minutes and then incubated with secondary antibodies diluted in PBS overnight at 4°C. The next day, samples were washed 3 times with PBS and incubated with Hoechst for 20 minutes to stain DNA/nuclei. Primary antibodies were chicken anti-GFP (Thermo Fisher Scientific), Rabbit anti-G6P (Boster Bio), Mouse Anti-CNX99A (DSHB Biology), Rabbit Anti-GM130 (abcam-ab30637). For DNA staining, either DAPI or Hoechst was used. The secondary antibodies that were used in this study were anti-chicken Alexa488 (Thermo Fisher Scientific), anti-rabbit Alexa Fluor 647 (abcam, ab150075), and anti-mouse Alexa 488 (abcam, ab150105). The polyclonal antibody against *Drosophila* G6Pase was generated using the peptide VREAFKWCPEPTTYLR (amino acid 248-263; Boster Bio).

### Sperm tail-beat frequency analysis

For tailbeat frequency measurements, sperm was isolated from seminal vesicles 2 days after eclosion (males) or spermathecae (female) 1 day after mating, via dissection of reproductive organs in PBS. Reproductive organs were sliced open, and sperm was released into 20μl of PBS on a glass slide. Sperm motility video micrographs were taken on an Olympus FV3000 scanning confocal microscope using an Olympus 20x/0.75NA UPLSAPO air objective lens. Images were acquired by Olympus Fluoview Software with the transmitted detector in resonant scanning mode, imaging at 0.067 μSec/px, and pixel size of 0.211 µm x 0.211 µm. The total ROI was set to include the whole sperm tail and was aimed to be kept similar in dimension. 1000 frames were collected per image. 640 nm laser was used as the light source with a laser power of 0.55% and transmitted detector sensitivity of 272V. DIC prism and polarizer were included in the light path.

To measure sperm tail-beat frequency, .Oir time series were saved as .Avi video clips. The frame per second (fps) was set, such as the number of frames divided by the length of the video. To measure the tail beat frequency, we followed the protocol by Hafezi and colleagues (Hafezi et al., 2024). Briefly, videos were analyzed using the ffmpeg plugin of FIJI (RRID:SCR_002285). The tail beat frequency of the fastest-beating sperm was measured by generating the kymograph. The number of beats and frames were counted for the sperm tail of interest. Beat frequency was measured using the below formula:

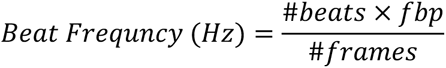

### Sperm Count in testes

Virgin males were separated after eclosion and kept isolated for 3 to 4 days, and the number of sperm in the seminal vesicle was counted using Imaris 10.1.1 software (RRID:SCR_007370) following the protocol of Hafezi and collaborators (Hafezi et al., 2024), with minor modifications. Briefly, testes were dissected, fixed, and stained for the nuclei using Hoechst. The entire seminal vesicle area were scanned, and accessory glands were excluded. Using the “Surfaces” function, sperm heads were detected using 1μm surface grain and sphericity set to less than 0.6).

### Sperm Transfer

To assess sperm transfer after the mating was conducted as described by Köttgen et al. (Köttgen et al., 2011) with minor modifications. Briefly, virgin *dj-GFP*/*dj-GFP* (control) males and *dj-GFP*/*dj-GFP*; *G6P^MIC^/G6P^MIC^* (experimental) males were kept separated for 24 to 36 hours before mating was set up to *w^1118^* virgin females. Immediately after the mating, females were separated, and their lower reproductive tract was dissected in PBS. The ovaries and digestive tract were removed carefully. The reproductive tracts of females were transferred to a glass slide with 20μl of PBS. Real-time live imaging of fluorescently labeled sperm in females was performed using a Nikon T1 A1R inverted confocal microscope. Microscopic analysis was performed within two hour (0 days), 5, 10 and 15 days after mating termination for 10 females. Sperm entry into the eggs was assessed for virgin males (*dj-GFP* in a wild type and *G6P^MIC/MIC^* background). Males and females were left to mate for 24h, and after that, the females were left on grape juice agar plates to lay eggs for 1h. Every hour the flies were flipped and number of eggs were counted on the plate and they were transferred to the glass slide and checked for the presence of sperm in the embryo. The numbers were presented as percentages of GFP positive on each plate.

### Seahorse Analysis

We used method described by Neville and colleagues to measure whole testes metabolism, with minor modifications (Neville et al., 2018). The testes were dissected as described for RNA extraction to measure the glycolytic rate. The software measures the extracellular acidification rates of the media. 96 well Seahorse plates were twice coated with Cell-Tak (50μg/mL, Corning® Cell-Tak^TM^ Cell and Tissue Adhesive, Cat. No. 354249). Testes were placed in the middle of the well and pushed down by forceps. To measure glycolytic rate proton efflux, we used the base medium without Phenol (Agilent, 103335-100), supplemented with HEPES (Agilent, 103337-100), 2mM Glutamine, 1mM sodium pyruvate, and 10mM glucose. Injection port A was used to supply 20μL of 50μM rotenone and antimycin-A. Injection port B was used to supply 22uL of 1M, 2-Deoxyglutarate. Analysis of glycolytic rate was done following Agilent guidelines (Agilent Technologies, Seahorse XF Glycolysis Stress Test Kit User Guide). After the assay, tissue was homogenized and lysed using default lysis buffer (50 mM Tris (pH 7.5), 125 mM NaCl, 5% glycerol, 0.2% IGEPAL, 1.5 mM MgCl_2_, 1 mM DTT, 25 mM NaF, 1 mM Na_3_VO_4_, 1 mM EDTA and 2 × Complete protease inhibitor (Roche, Indianapolis, IN) on ice for 10 min. The plate was then centrifuged, and the supernatant was collected.

### Measurement of bulk ATP

Total ATP levels in testes were measured using the ATPlite luminescence assay system (PerkinElmer), a luciferase-based assay. Measurements were carried out as described by Nellas and colleagues (Nellas et al., 2022). *Drosophila* testes were dissected in PBS and transferred to the wells of flat bottom polystyrene assay plates (Costar® 3917) containing 100 ml of PBS (1 pair/well). Samples were lysed by adding 50 ml of cell lysis solution and shaking at 700 rpm for 10 minutes. 50ml of luciferase substrate solution was added, shaking at 700 rpm for 5 minutes. After incubating in the dark for 10 minutes, luciferase activity was measured using a PerkinElmer EnVision plate reader. ATP concentration was calculated using a standard curve, generated through serial dilution of a 10 mM ATP stock solution.

## REFERENCES

Banka, S., Chervinsky, E., Newman, W.G., Crow, Y.J., Yeganeh, S., Yacobovich, J., Donnai, D., Shalev, S., 2011. Further delineation of the phenotype of severe congenital neutropenia type 4 due to mutations in G6PC3. Eur J Hum Genet 19, 18–22. 10.1038/ejhg.2010.136

Bertolotti, A., Zhang, Y., Hendershot, L.M., Harding, H.P., Ron, D., 2000. Dynamic interaction of BiP and ER stress transducers in the unfolded-protein response. Nat Cell Biol 2, 326–332. 10.1038/35014014

Bommer, G.T., Van Schaftingen, E., Veiga-da-Cunha, M., 2020. Metabolite Repair Enzymes Control Metabolic Damage in Glycolysis. Trends Biochem Sci 45, 228–243. 10.1016/j.tibs.2019.07.004

Bosma, K.J., Rahim, M., Oeser, J.K., McGuinness, O.P., Young, J.D., O’Brien, R.M., 2020a. G6PC2 confers protection against hypoglycemia upon ketogenic diet feeding and prolonged fasting. Mol Metab 41, 101043. 10.1016/j.molmet.2020.101043

Bosma, K.J., Rahim, M., Singh, K., Goleva, S.B., Wall, M.L., Xia, J., Syring, K.E., Oeser, J.K., Poffenberger, G., McGuinness, O.P., Means, A.L., Powers, A.C., Li, W.-H., Davis, L.K., Young, J.D., O’Brien, R.M., 2020b. Pancreatic islet beta cell-specific deletion of G6pc2 reduces fasting blood glucose. J Mol Endocrinol 64, 235–248. 10.1530/JME-20-0031

Boztug, K., Klein, C., 2009. Novel genetic etiologies of severe congenital neutropenia. Curr Opin Immunol 21, 472–480. 10.1016/j.coi.2009.09.003

Burchell, A., Allan, B.B., Hume, R., 1994. Glucose-6-phosphatase proteins of the endoplasmic reticulum. Mol Membr Biol 11, 217–227. 10.3109/09687689409160431

Cairrão, F., Santos, C.C., Le Thomas, A., Marsters, S., Ashkenazi, A., Domingos, P.M., 2022. Pumilio protects Xbp1 mRNA from regulated Ire1-dependent decay. Nat Commun 13, 1587. 10.1038/s41467-022-29105-x

Chou, J.Y., Jun, H.S., Mansfield, B.C., 2015. Type I glycogen storage diseases: disorders of the glucose-6-phosphatase/glucose-6-phosphate transporter complexes. J Inherit Metab Dis 38, 511–519. 10.1007/s10545-014-9772-x

Coe, H., Bedard, K., Groenendyk, J., Jung, J., Michalak, M., 2008. Endoplasmic reticulum stress in the absence of calnexin. Cell Stress Chaperones 13, 497–507. 10.1007/s12192-008-0049-x

Cytological analysis of spermatogenesis: live and fixed preparations of Drosophila testes., 2014.. United States. 10.3791/51058

Danilczyk, U.G., Cohen-Doyle, M.F., Williams, D.B., 2000. Functional relationship between calreticulin, calnexin, and the endoplasmic reticulum luminal domain of calnexin. J Biol Chem 275, 13089–13097. 10.1074/jbc.275.17.13089

Dansereau, D.A., Lasko, P., 2008. The development of germline stem cells in Drosophila. Methods Mol Biol 450, 3–26. 10.1007/978-1-60327-214-8_1

Dienel, G.A., 2020. Hypothesis: A Novel Neuroprotective Role for Glucose-6-phosphatase (G6PC3) in Brain-To Maintain Energy-Dependent Functions Including Cognitive Processes. Neurochem Res 45, 2529–2552. 10.1007/s11064-020-03113-z

Fabian, L., Brill, J.A., 2012. Drosophila spermiogenesis: Big things come from little packages. Spermatogenesis 2, 197–212. 10.4161/spmg.21798

Fitch, K.R., Wakimoto, B.T., 1998. The paternal effect gene ms(3)sneaky is required for sperm activation and the initiation of embryogenesis in Drosophila melanogaster. Dev Biol 197, 270–282. 10.1006/dbio.1997.8852

Fuller, M.T., 1998. Genetic control of cell proliferation and differentiation in Drosophila spermatogenesis. Semin Cell Dev Biol 9, 433–444. 10.1006/scdb.1998.0227

Hafezi, Y., Omurzakov, A., Carlisle, J.A., Caldas, I.V., Wolfner, M.F., Clark, A.G., 2024. The Drosophila melanogaster Y-linked gene, WDY, is required for sperm to swim in the female reproductive tract. Commun Biol 7, 90. 10.1038/s42003-023-05717-x

Hayee, B., Antonopoulos, A., Murphy, E.J., Rahman, F.Z., Sewell, G., Smith, B.N., McCartney, S., Furman, M., Hall, G., Bloom, S.L., Haslam, S.M., Morris, H.R., Boztug, K., Klein, C., Winchester, B., Pick, E., Linch, D.C., Gale, R.E., Smith, A.M., Dell, A., Segal, A.W., 2011. G6PC3 mutations are associated with a major defect of glycosylation: a novel mechanism for neutrophil dysfunction. Glycobiology 21, 914–924. 10.1093/glycob/cwr023

Intra, J., Cenni, F., Perotti, M.-E., 2006. An alpha-L-fucosidase potentially involved in fertilization is present on Drosophila spermatozoa surface. Mol Reprod Dev 73, 1149–1158. 10.1002/mrd.20425

Jackson, M.R., Cohen-Doyle, M.F., Peterson, P.A., Williams, D.B., 1994. Regulation of MHC class I transport by the molecular chaperone, calnexin (p88, IP90). Science 263, 384–387. 10.1126/science.8278813

Jun, H.S., Lee, Y.M., Cheung, Y.Y., McDermott, D.H., Murphy, P.M., De Ravin, S.S., Mansfield, B.C., Chou, J.Y., 2010. Lack of glucose recycling between endoplasmic reticulum and cytoplasm underlies cellular dysfunction in glucose-6-phosphatase-beta-deficient neutrophils in a congenital neutropenia syndrome. Blood 116, 2783–2792. 10.1182/blood-2009-12-258491

Jun, H.S., Lee, Y.M., Song, K.D., Mansfield, B.C., Chou, J.Y., 2011. G-CSF improves murine G6PC3-deficient neutrophil function by modulating apoptosis and energy homeostasis. Blood 117, 3881–3892. 10.1182/blood-2010-08-302059

Köttgen, M., Hofherr, A., Li, W., Chu, K., Cook, S., Montell, C., Watnick, T., 2011. Drosophila sperm swim backwards in the female reproductive tract and are activated via TRPP2 ion channels. PLoS One 6, e20031. 10.1371/journal.pone.0020031

Li, H., Janssens, J., De Waegeneer, M., Kolluru, S.S., Davie, K., Gardeux, V., Saelens, W., David, F.P.A., Brbić, M., Spanier, K., Leskovec, J., McLaughlin, C.N., Xie, Q., Jones, R.C., Brueckner, K., Shim, J., Tattikota, S.G., Schnorrer, F., Rust, K., Nystul, T.G., Carvalho-Santos, Z., Ribeiro, C., Pal, S., Mahadevaraju, S., Przytycka, T.M., Allen, A.M., Goodwin, S.F., Berry, C.W., Fuller, M.T., White-Cooper, H., Matunis, E.L., DiNardo, S., Galenza, A., O’Brien, L.E., Dow, J.A.T., Jasper, H., Oliver, B., Perrimon, N., Deplancke, B., Quake, S.R., Luo, L., Aerts, S., Agarwal, D., Ahmed-Braimah, Y., Arbeitman, M., Ariss, M.M., Augsburger, J., Ayush, K., Baker, C.C., Banisch, T., Birker, K., Bodmer, R., Bolival, B., Brantley, S.E., Brill, J.A., Brown, N.C., Buehner, N.A., Cai, X.T., Cardoso-Figueiredo, R., Casares, F., Chang, A., Clandinin, T.R., Crasta, S., Desplan, C., Detweiler, A.M., Dhakan, D.B., Donà, E., Engert, S., Floc’hlay, S., George, N., González-Segarra, A.J., Groves, A.K., Gumbin, S., Guo, Y., Harris, D.E., Heifetz, Y., Holtz, S.L., Horns, F., Hudry, B., Hung, R.-J., Jan, Y.N., Jaszczak, J.S., Jefferis, G.S.X.E., Karkanias, J., Karr, T.L., Katheder, N.S., Kezos, J., Kim, A.A., Kim, S.K., Kockel, L., Konstantinides, N., Kornberg, T.B., Krause, H.M., Labott, A.T., Laturney, M., Lehmann, R., Leinwand, S., Li, J., Li, J.S.S., Li, Kai, Li, Ke, Li, L., Li, T., Litovchenko, M., Liu, H.-H., Liu, Y., Lu, T.-C., Manning, J., Mase, A., Matera-Vatnick, M., Matias, N.R., McDonough-Goldstein, C.E., McGeever, A., McLachlan, A.D., Moreno-Roman, P., Neff, N., Neville, M., Ngo, S., Nielsen, T., O’Brien, C.E., Osumi-Sutherland, D., Özel, M.N., Papatheodorou, I., Petkovic, M., Pilgrim, C., Pisco, A.O., Reisenman, C., Sanders, E.N., Dos Santos, G., Scott, K., Sherlekar, A., Shiu, P., Sims, D., Sit, R.V., Slaidina, M., Smith, H.E., Sterne, G., Su, Y.-H., Sutton, D., Tamayo, M., Tan, M., Tastekin, I., Treiber, C., Vacek, D., Vogler, G., Waddell, S., Wang, W., Wilson, R.I., Wolfner, M.F., Wong, Y.-C.E., Xie, A., Xu, J., Yamamoto, S., Yan, J., Yao, Z., Yoda, K., Zhu, R., Zinzen, R.P., 2022. Fly Cell Atlas: A single-nucleus transcriptomic atlas of the adult fruit fly. Science 375, eabk2432. 10.1126/science.abk2432

Lindsey, D., L., Tokuyashu, K.T., 1980. Spermatogenesis, in: Genetics and Biology of Drosophila. Academic Press, New York, pp. 225–294.

Loppin, B., Dubruille, R., Horard, B., 2015. The intimate genetics of Drosophila fertilization. Open Biol 5. 10.1098/rsob.150076

Matsuda, H., Yamada, T., Yoshida, M., Nishimura, T., 2015. Flies without trehalose. J Biol Chem 290, 1244–1255. 10.1074/jbc.M114.619411

McCullough, E.L., Whittington, E., Singh, A., Pitnick, S., Wolfner, M.F., Dorus, S., 2022. The life history of Drosophila sperm involves molecular continuity between male and female reproductive tracts. Proc Natl Acad Sci U S A 119, e2119899119. 10.1073/pnas.2119899119

Miyamoto, T., Amrein, H., 2019. Neuronal Gluconeogenesis Regulates Systemic Glucose Homeostasis in Drosophila melanogaster. Curr Biol 29, 1263–1272.e5. 10.1016/j.cub.2019.02.053

Miyamoto, T., Amrein, H., 2017. Gluconeogenesis: An ancient biochemical pathway with a new twist. Fly (Austin) 11, 218–223. 10.1080/19336934.2017.1283081

Miyamoto, T., Hedjazi, S., Miyamoto, C., Amrein, H., 2024. Drosophila Neuronal Glucose 6 Phosphatase is a modulator of Neuropeptide Release that regulates muscle glycogen stores via FMRFamide signaling. Proceedings of the National Academy of Sciences of the United States of America 2023.11.28.568950. 10.1101/2023.11.28.568950

Morciano, P., Di Giorgio, M.L., Tullo, L., Cenci, G., 2021. The Organization of the Golgi Structures during Drosophila Male Meiosis Requires the Citrate Lyase ATPCL. Int J Mol Sci 22. 10.3390/ijms22115745

Nellas, I., Iyer, K.V., Iglesias-Artola, J.M., Pippel, M., Nadler, A., Eaton, S., Dye, N.A., 2022. Hedgehog signaling can enhance glycolytic ATP production in the Drosophila wing disc. EMBO Rep 23, e54025. 10.15252/embr.202154025

Neville, K.E., Bosse, T.L., Klekos, M., Mills, J.F., Weicksel, S.E., Waters, J.S., Tipping, M., 2018. A novel ex vivo method for measuring whole brain metabolism in model systems. J Neurosci Methods 296, 32–43. 10.1016/j.jneumeth.2017.12.020

Okabe, M., 2018. Sperm-egg interaction and fertilization: past, present, and future. Biol Reprod 99, 134–146. 10.1093/biolre/ioy028

Perotti, M.E., Cattaneo, F., Pasini, M.E., Vernì, F., Hackstein, J.H., 2001. Male sterile mutant casanova gives clues to mechanisms of sperm-egg interactions in Drosophila melanogaster. Mol Reprod Dev 60, 248–259. 10.1002/mrd.1085

Perotti, M.E., Pasini, M.E., 1995. Glycoconjugates of the surface of the spermatozoa of Drosophila melanogaster: a qualitative and quantitative study. J Exp Zool 272, 311–318. 10.1002/jez.1402720409

Perotti, M.E., Riva, A., 1988. Concanavalin A binding sites on the surface of Drosophila melanogaster sperm: a fluorescence and ultrastructural study. J Ultrastruct Mol Struct Res 100, 173–182. 10.1016/0889-1605(88)90024-9

Santel, A., Winhauer, T., Blümer, N., Renkawitz-Pohl, R., 1997. The Drosophila don juan (dj) gene encodes a novel sperm specific protein component characterized by an unusual domain of a repetitive amino acid motif. Mech Dev 64, 19–30. 10.1016/s0925-4773(97)00031-2

Uemura, A., Oku, M., Mori, K., Yoshida, H., 2009. Unconventional splicing of XBP1 mRNA occurs in the cytoplasm during the mammalian unfolded protein response. J Cell Sci 122, 2877– 2886. 10.1242/jcs.040584

Veiga-da-Cunha, M., Chevalier, N., Stephenne, X., Defour, J.-P., Paczia, N., Ferster, A., Achouri, Y., Dewulf, J.P., Linster, C.L., Bommer, G.T., Van Schaftingen, E., 2019. Failure to eliminate a phosphorylated glucose analog leads to neutropenia in patients with G6PT and G6PC3 deficiency. Proc Natl Acad Sci U S A 116, 1241–1250. 10.1073/pnas.1816143116

Veiga-da-Cunha, M., Van Schaftingen, E., Bommer, G.T., 2020. Inborn errors of metabolite repair. J Inherit Metab Dis 43, 14–24. 10.1002/jimd.12187

Wakimoto, B.T., Lindsley, D.L., Herrera, C., 2004. Toward a comprehensive genetic analysis of male fertility in Drosophila melanogaster. Genetics 167, 207–216. 10.1534/genetics.167.1.207

Wetzker, C., Froschauer, C., Massino, C., Reinhardt, K., 2024. Drosophila melanogaster sperm turn more oxidative in the female. J Exp Biol 227. 10.1242/jeb.247775

Wigby, S., Brown, N.C., Allen, S.E., Misra, S., Sitnik, J.L., Sepil, I., Clark, A.G., Wolfner, M.F., 2020. The Drosophila seminal proteome and its role in postcopulatory sexual selection. Philos Trans R Soc Lond B Biol Sci 375, 20200072. 10.1098/rstb.2020.0072

Wilson, K.L., Fitch, K.R., Bafus, B.T., Wakimoto, B.T., 2006. Sperm plasma membrane breakdown during Drosophila fertilization requires sneaky, an acrosomal membrane protein. Development 133, 4871–4879. 10.1242/dev.02671

Yeshayahu, Y., Asaf, R., Dubnov-Raz, G., Schiby, G., Simon, A.J., Lev, A., Somech, R., 2014. Testicular failure in a patient with G6PC3 deficiency. Pediatr Res 76, 197–201. 10.1038/pr.2014.64

Zelinger, E., Brumfeld, V., Rechav, K., Waiger, D., Kossovsky, T., Heifetz, Y., 2024. Three-dimensional correlative microscopy of the Drosophila female reproductive tract reveals modes of communication in seminal receptacle sperm storage. Commun Biol 7, 155. 10.1038/s42003-024-05829-y

