## Supplemental Figures for "Glucose-6-phosphatase is required for organelle reorganization, energy metabolism and motility of *Drosophila* sperm"

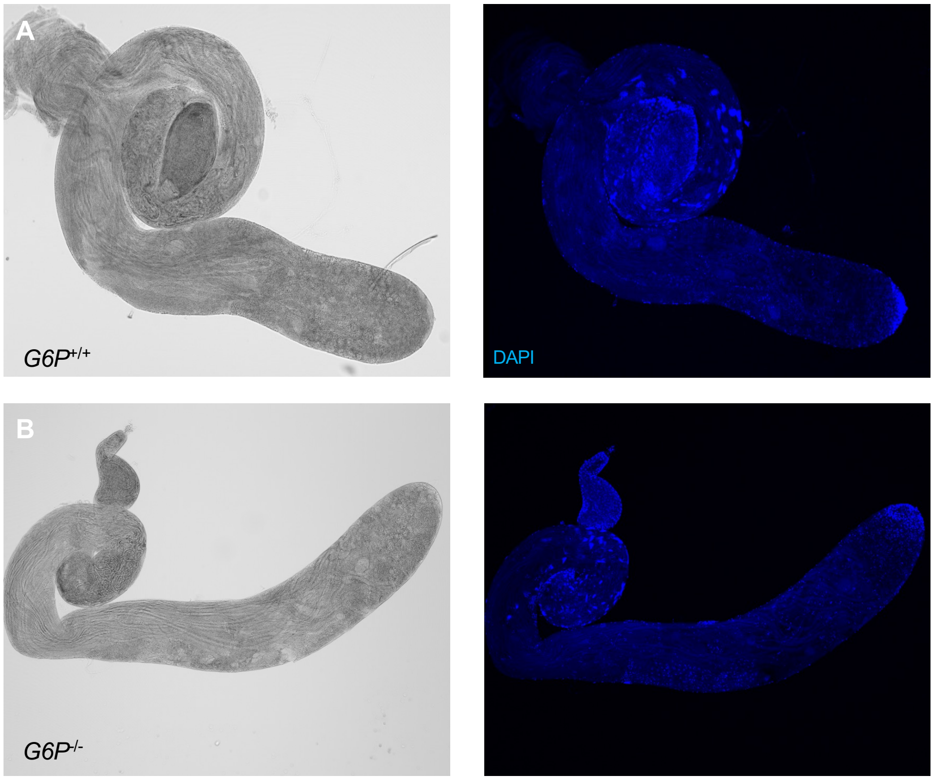


*G6P* mutant males show no apparent defects in spermatogenesis:  A testis of a wild type (top) and *G6P* mutant male (bottom) shows normal progression through spermatogenesis germ stem cells at the apical tip and testis filled with mature sperm

**Supp Figure 1**

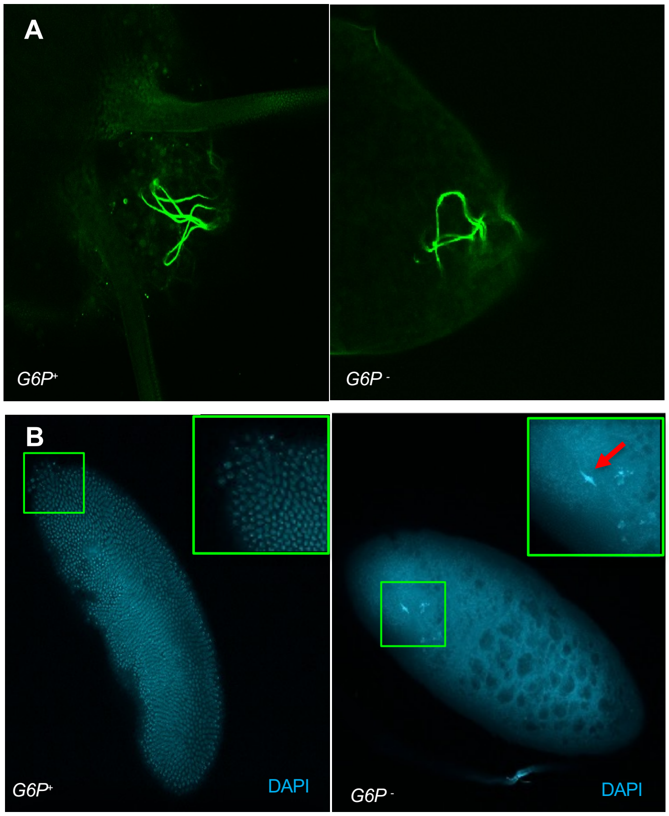


Top: Wild type (left) and *G6P-* (right) sperm containing a *dj-GFP* transgene, visualized using live imaging in a fertilized embryo. GFP-labeled sperm tail is seen in 85% of eggs of females mated to wild type males, but only in about 8% of eggs of males mated to *G6P* mutant males (See Figure 4I).

Bottom: DAPI staining of an embryo/egg about 3 hours after egg laying. The embryo fertilized by wild type sperm stared to develop and at the cellular blastoderm stage, while the egg fertilized by the *G6P-* sperm failed to develop. Failure to break down the sperm membrane is apparent by the elongated male nuclear stain of DNA (red arrow).

**Supp Figure 2**

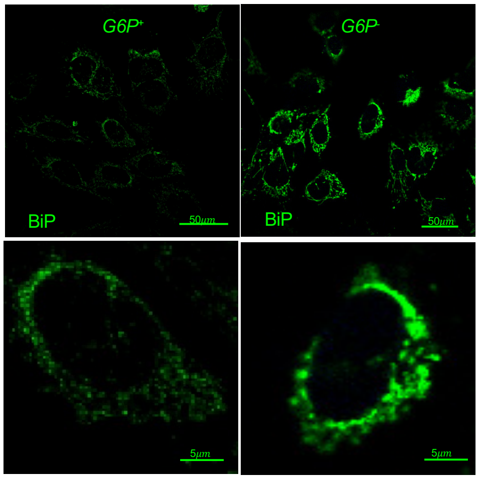


Upregulation of Binding immunoglobulin Protein in *G6P* mutant spermatocytes (left), visualized by anti-BiP antibody staining. Same microscope setting were used to allow comparison of immunoreactivity between the two genotypes. Images below show an enlarged spermatocytes of the view shown at the top.

**Supp Figure 3**

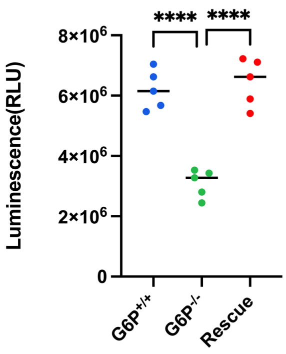


Total ATP levels in testes of *G6P* mutant (green), wild type (blue) and rescue control males (*Y/yw; G6P-/G6P-; GenG6P rescue/+,* red) males was measured using the ATPlite kit from PerkinElmer.

**Supp Figure 4**

GlyP (CG7254)

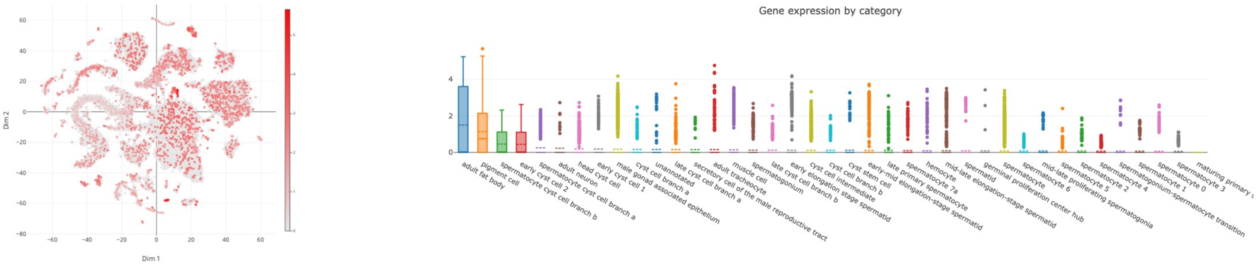


Agl (CG9485)

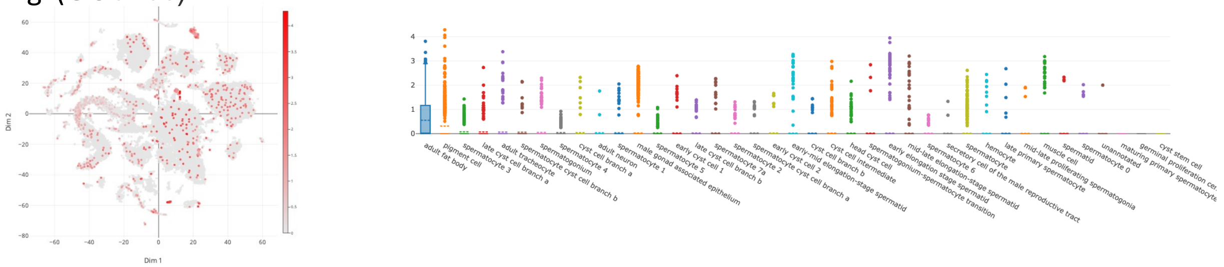


GlyS (CG6904)

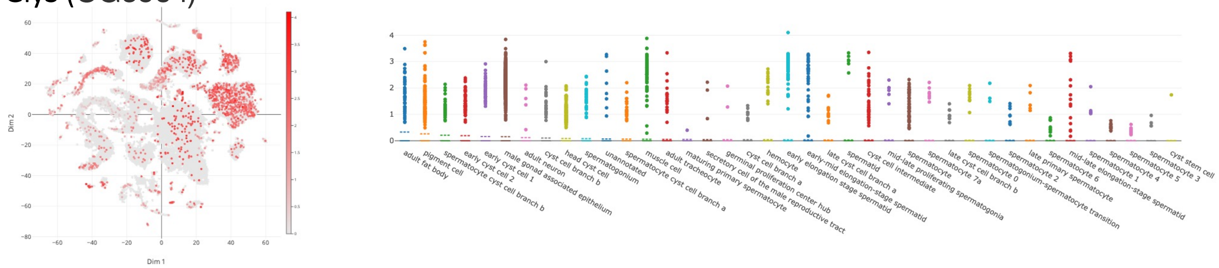


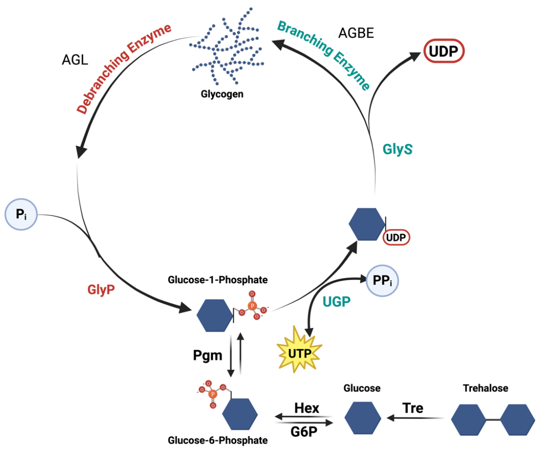


Genes encoding key enzymes involved in glycogen metabolism are not expressed in spermatocytes. Tissues/cells are listed based on gene expression level of indicted genes from left (high) to right (low).

**Supp Figure 5**

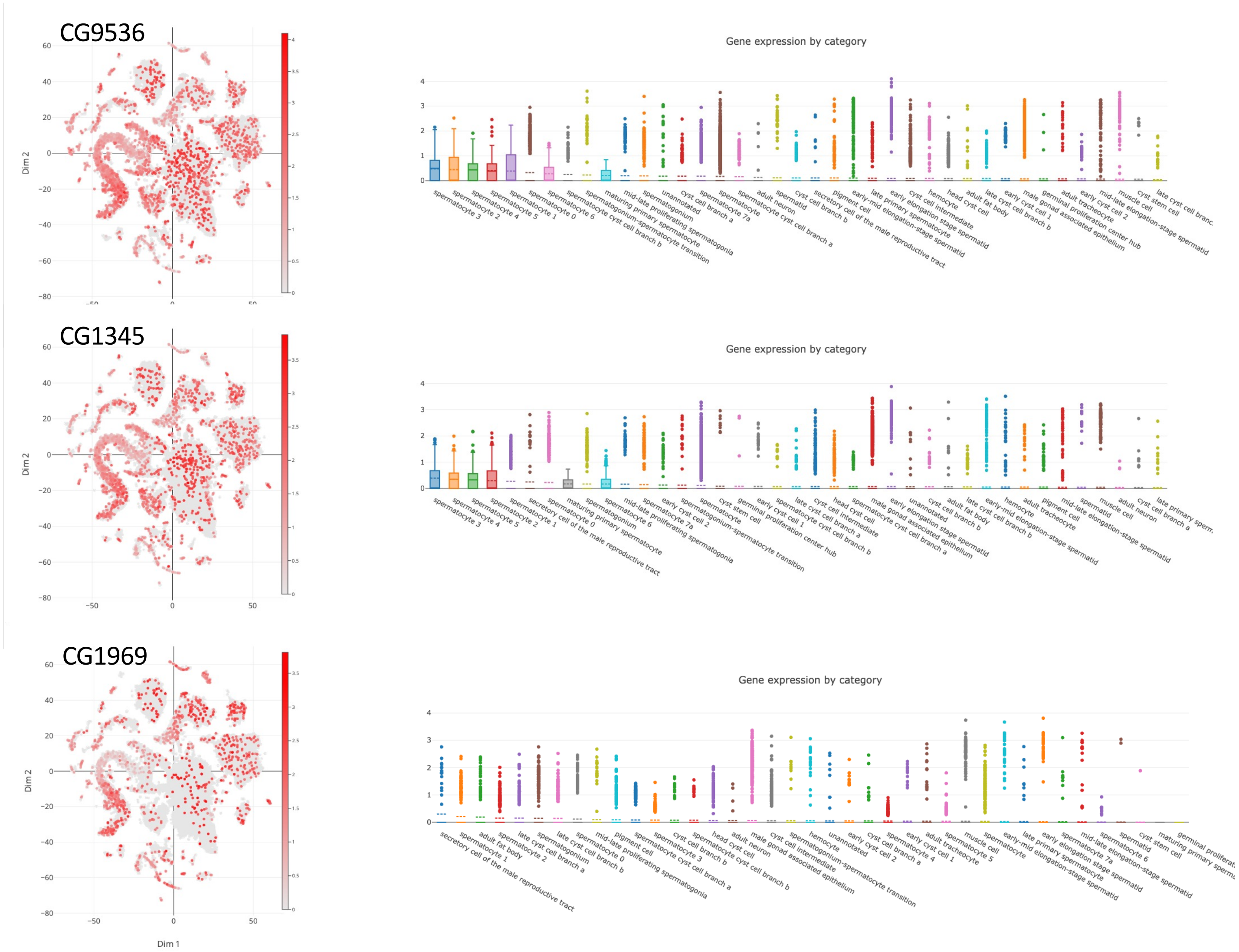


Key glycosylation enzymes, such as the fly homolog of mammalian TMEM 115 (CG9536), Glutamine:fructose-6-phosphate aminotransferase 2 (CG1345) and Glucosamine-phosphate N-acetyltransferase (CG1969) are mainly or specifically expressed in spermatocytes. Tissues/cells are listed based on gene expression level of indicted genes from left (high) to right (low).

**Supp Figure 6**

### Fertility rates

| | $G6P^{+/+}$ | | $G6P^{-/-}$ | | G6P_rescue | |
|---|---|---|---|---|---|---|
| | E laid | L hatched | E laid | L hatched | E laid | L hatched |
| 1 | 18 | 15 | 20 | 0 | 22 | 19 |
| 2 | 21 | 17 | 21 | 1 | 20 | 16 |
| 3 | 25 | 23 | 22 | 0 | 19 | 18 |
| 4 | 21 | 20 | 25 | 0 | 18 | 15 |
| 5 | 20 | 18 | 19 | 2 | 18 | 16 |
| 6 | 24 | 20 | 25 | 0 | 25 | 20 |
| 7 | 23 | 20 | 23 | 0 | 23 | 20 |
| 8 | 21 | 19 | 20 | 1 | 20 | 12 |
| 9 | 19 | 15 | 22 | 0 | 21 | 19 |
| 10 | 27 | 21 | 20 | 0 | 25 | 16 |
| 11 | 21 | 20 | 20 | 0 | 19 | 17 |
| 12 | 25 | 20 | 21 | 0 | 21 | 17 |
| 13 | 23 | 20 | 25 | 1 | 21 | 16 |
| 14 | 24 | 19 | 19 | 0 | 20 | 14 |
| 15 | 22 | 19 | 19 | 0 | 19 | 15 |
| 16 | 20 | 17 | 20 | 0 | 28 | 24 |
| 17 | 21 | 16 | 18 | 0 | 22 | 19 |
| 18 | 19 | 15 | 20 | 0 | 22 | 18 |
| 19 | 19 | 15 | 23 | 0 | 20 | 15 |
| 20 | 23 | 20 | 20 | 2 | 20 | 15 |
| average | 21.8 | 18.45 | 21.1 | 0.35 | 21,15 | 17.05 |
| **Hatching rate(%)** | **84.63** | | **1.66** | | **80.61** | |

**Supp Table 1**
